# Keratin 8 is a scaffolding and regulatory protein of ERAD complexes

**DOI:** 10.1101/2022.02.01.478623

**Authors:** Iwona M. Pranke, Benoit Chevalier, Aiswarya Premchandar, Nesrine Baatallah, Kamil F. Tomaszewski, Sara Bitam, Danielle Tondelier, Anita Golec, Jan Stolk, Gergely L. Lukacs, Pieter S. Hiemstra, Michal Dadlez, David A. Lomas, James A. Irving, Agnes Delaunay-Moisan, Eelco van Anken, Alexandre Hinzpeter, Isabelle Sermet-Gaudelus, Aleksander Edelman

## Abstract

Early recognition and enhanced degradation of misfolded proteins by the endoplasmic reticulum (ER) quality control and ER-associated degradation (ERAD) cause defective protein secretion and membrane targeting, as exemplified for Z-alpha 1 antitrypsin (Z-A1AT), responsible for alpha-1-antitrypsin deficiency (A1ATD) and F508del-CFTR (cystic fibrosis transmembrane conductance regulator) responsible for cystic fibrosis (CF).

Prompted by our previous observation that decreasing Keratin 8 (K8) expression increased trafficking of F508del-CFTR to the plasma membrane, we investigated whether K8 impacts trafficking of soluble misfolded Z-A1AT protein. The subsequent goal of this study was to elucidate the mechanism underlying the K8-dependent regulation of protein trafficking, focusing on the ERAD pathway.

The results show that diminishing K8 concentration in HeLa cells enhances secretion of both Z-A1AT and wild type (WT) A1AT with a 13-fold and 4-fold increase, respectively. K8 down-regulation triggers ER failure and cellular apoptosis when ER stress is jointly elicited by conditional expression of the μ_s_ heavy chains, as previously shown for Hrd1 knock-out. Simultaneous K8 silencing and Hrd1 knock-out did not show any synergistic effect, consistent with K8 acting in the Hrd1-governed ERAD step. Fractionation experiments reveal that K8 is recruited to ERAD complexes containing Derlin2, Sel1 and Hrd1 proteins upon expression of Z/WT-A1AT and F508del-CFTR. Treatment of the cells with c407, a small molecule inhibiting K8 interaction, decreases K8 and Derlin2 recruitment to high-order ERAD complexes. This was associated with increased Z-A1AT secretion in both HeLa and Z-homozygous A1ATD patients’ respiratory cells. Overall, we provide evidence that K8 acts as an ERAD modulator. It may play a scaffolding protein role for early-stage ERAD complexes, regulating Hrd1-governed retrotranslocation initiation/ubiquitination processes. Targeting K8-containing ERAD complexes is an attractive strategy for the pharmacotherapy of A1ATD.

## Introduction

Proper protein folding represents one of the primary challenges for cells to maintain proteostasis. This complex process is challenged by the intrinsic properties of proteins or mutations in protein-coding genes. Proteins entering the secretion pathway that fail to fold, assemble or post-translationally mature properly are eliminated by ERAD. ERAD relies on multiple dynamic protein complexes acting sequentially to recognise, tether, ubiquitin-tag and extract misfolded substrates from the ER, thereby enabling their proteasomal degradation in the cytosol (Christianson and Ye, 2014) (Mehrtash and Hochstrasser, 2019) (Ruggiano et al., 2014). Distinct ERAD pathways accommodate the variety of topologies and post-translational modifications encountered in misfolded protein substrates. The partial redundancy of ERAD factors allows cells to oversee proteostatic stress and maintain ER homeostasis coordinately. ERAD pathways can be divided into the following functional modules: (i) substrate recognition;(ii) retro-translocation initiation with substrate unfolding involving mainly multifunctional Derlin proteins; (iii) ubiquitination of the partially dislocated substrate, assisted by different membrane-bound E3 ubiquitin ligases (e.g. a multimeric complex, containing Hrd1 E3 ligase coupled to substrate recognition protein Sel1) and associated ubiquitin-conjugating enzymes E2s according to substrates (Baldridge and Rapoport, 2016) (Stein et al., 2014) (Vasic et al., 2020); therefore licensing (iv) the dislocation process ensured by p97/VCP ATP-ase providing a pulling force (Garza et al., 2009) (Nakatsukasa et al., 2008) (Rabinovich et al., 2002); and finally (v) substrate degradation by 26S proteasome.

Protein misfolding underlies two frequent pulmonary genetic diseases, A1ATD and CF. The most common disease-causing protein folding mutations are Glu342Lys (Z) in A1AT and F508del in CFTR (Stoller and Aboussouan, 2012) (Ghouse et al., 2014) (Pranke et al., 2019). Over 99% of F508del-CFTR and ∼70% Z-A1AT are degraded by ERAD (Jensen et al., 1995) (Teckman et al., 2001) (Kroeger et al., 2009a). Both misfolded proteins form complexes with ER-chaperones (e.g., Calnexin) (Kim and Skach, 2012) (Schmidt and Perlmutter, 2005), ubiquitin ligases (e.g., Hrd1 and its partner Sel1) (Lomas et al., 1992) (Khodayari et al., 2017a) (Glenn et al., 2012) (Shen et al., 2006) (Graham et al., 1990), and dislocating protein p97/VCP (Perlmutter, 2006) (Khodayari et al., 2017b) (Carlson et al., 2006). Consequently, misfolded F508del-CFTR is less efficiently targeted to the apical membrane (PM) of epithelial cells and secretion of misfolded Z-A1AT is strongly decreased (to ∼10–15% of WT level). In addition, the folding efficacy of WT-CFTR is limited (Okiyoneda and Lukacs, 2012) (Okiyoneda et al., 2013), and the thermodynamic state of WT-A1AT is suboptimal (Chakraborty and Teckman, 2014), both features favouring ERAD degradation of the wild type isotypes (Varga et al., 2004). Additionally, for Z-A1AT, which is highly expressed in the hepatocyte, its accumulation in the ER also activates classic autophagy (Kroeger et al., 2009a) and ER-to-lysosome-associated degradation (Fregno et al., 2018).

An increasing body of evidence suggests that Keratin 8 (K8) and Keratin 18 (K18), proteins forming intermediary filament (IF) heterodimers in simple epithelia (Herrmann and Aebi, 2004), can regulate protein targeting to the apical plasma membrane (Coulombe and Omary, 2002) (Toivola et al., 2004) (Mashukova et al., 2016). They also display additional roles beyond mechanical and structural functions, including protein polyubiquitination in a pro-inflammatory context (Dong et al., 2016). Our previous studies (Davezac et al., 2004) (Colas et al., 2012a) revealed that K8 and K18 regulate F508del-CFTR trafficking. We have determined that K8 retained F508del-CFTR in the ER (Colas et al., 2012b), an effect that can be alleviated pharmacologically with small molecules, e.g., c407 (Odolczyk et al., 2013), and/or reducing K8 concentrations by siRNA (Colas et al., 2012a).

Here we show that K8 regulates the secretion of Z-A1AT. Since K8 affects the targeting of two ERAD substrates, Z-A1AT and F508del-CFTR, we challenged the hypothesis that K8 is part of the ERAD pathway. We gathered genetic and biochemical clues pointing to a new physiological function of K8 during the ERAD process. As for significant ERAD components (Hrd1, Sel1) (Vitale et al., 2019) in conditions triggering ER stress, we observed that K8 is essential to prevent ER failure and associated cell death. K8 co-sedimented with major ERAD components (Derlin2, Hrd1 and Sel1), consistent with K8 acting as a scaffold in the Hrd1-dependent ERAD pathway. Finally, we provide proof of concept that modulation of K8-containing ERAD complexes increases Z-A1AT secretion from HeLa, and patients derived primary human respiratory epithelial (HNE) cells and could be a target for pharmacotherapy in A1ATD.

### Results

### 1. Silencing K8 expression increases WT-A1AT and Z-A1AT secretion through the conventional secretory pathway

The implication of K8 in the secretion of WT-A1AT and misfolded Z-A1AT was evaluated in HeLa cells upon reduced K8 levels (shRNAK8, see Material and Methods section). Quantification of A1AT in the supernatants of control and shK8 cells showed significantly enhanced secretion of both Z-A1AT and WT-A1AT in shK8 cells (**Fig. 1A and B Ctrl and shCtrl**), with a 13-fold and 4-fold increase of secretion, respectively (**Fig. 1B Ctrl and shCtrl**). This secretion was associated with an increased intracellular pool of Z-A1AT (**Fig. 1C** and **D Ctrl and shCtrl**), suggesting its stabilisation.

**Figure 1.**
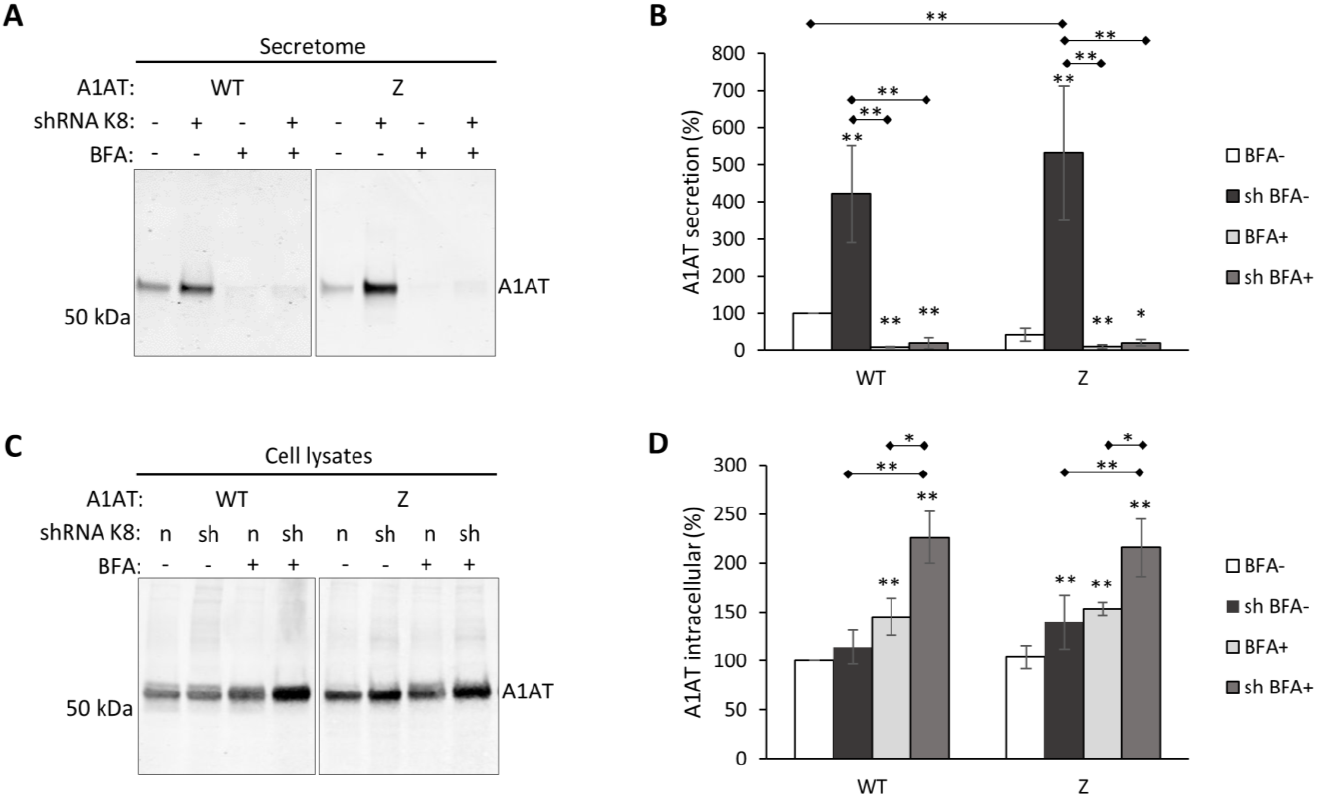
Silencing K8 in HeLa cells increases the secretion of Z-A1AT and WT-A1AT through conventional secretory pathway. **(A)** WB analysis of secretion of WT-A1AT/Z-A1AT in cell cultures with normal (shRNAK8-) and decreased (shRNAK8+) level of K8 expression after treatment with Brefeldin A 5μg/ml (BFA+) or vehicle (BFA-) for 7 hours. Representative image and quantification from n=9 independent experiments. (B) Quantification of secretion (mean±SD) normalised to total protein concentrations in respective cell lysates from at least four experiments.* < 0.05; ** < 0.005; *** < 0.0005 (Mann-Whitney test); * over the bar indicates p value vs. control (white boxes), if otherwise it is indicated. **(C)** WB analysis of intracellular WT-A1AT/Z-A1AT in cell cultures with normal (shRNAK8-) and decreased (shRNAK8+) level of K8 expression after treatment with Brefeldin A 5μg/ml (BFA+) or vehicle (BFA-) for 7 hours. Representative image and quantification from n=9 independent experiments. **(D)** Quantification of secretion (mean±SD) normalized to total protein concentrations in respective cell lysates from at least 4 experiments. *< 0.05; ** < 0.005; *** < 0.0005 (Mann-Whitney test); * over the bar indicates p value vs. control (white boxes), if otherwise it is indicated.

To determine if the observed increase in WT-/ Z-A1AT secretion was not an overall effect on proteins secretion, we performed three series of control experiments: (i) quantification of the secretome protein content did not differ in shK8 and K8-expressing (control) HeLa cells (**Supp. Fig. 2A**); (ii) secretion of transfected *Gaussia* luciferase was not modified (**Supp. Fig. 2B**) and, (iii) expression of endogenous Na^+^K^+^ATPase at the plasma membrane was not changed upon K8 silencing (**Supp. Fig. 2C**). Altogether these experiments suggest that K8-dependent regulation of secretion concerned misfolded Z-A1AT or unstable WT-A1AT.

The enhanced WT- and Z-A1AT secreted by shK8 cells had an identical SDS-gel migration pattern compared to WT- and Z-A1AT secreted by control cells (**Fig. 1A**, shRNAK8-), indicative of a similar glycosylation pattern and secretion occurring through the conventional secretory pathway. This was confirmed using Brefeldin A (BFA), an inhibitor of the conventional secretory pathway, which abolished WT-/Z-A1AT secretion from both control and shK8 cells (**Fig. 1A and B**). Upon BFA treatment, an increase of Z- and WT-A1AT intracellular concentrations was observed in shK8 cells compared to control cells (**Fig. 1C and D**), consistent with reduced degradation of WT-/Z-A1AT upon decreased K8 expression.

The accumulation of Z-A1AT upon BFA treatment in shK8 cells suggests an implication of K8 in the degradation *via* the ERAD or the autophagy pathways (Kroeger et al., 2009b). To test for a possible role of K8 silencing in rescuing Z-A1AT from autophagy, the effect of Wortmannin (1μM, 24h), an inhibitor of phosphoinositide 3-kinases, was tested. Intracellular accumulation of non-mature Z-A1AT treatment was observed without increased secretion (**Supp. Fig. 3A**). This suggests that autophagy was not involved in K8 silencing-dependent enhanced A1AT secretion. On the other hand, associated with the accumulation of Z-A1AT, an increased expression of Grp78/BiP, a marker of ER stress, was observed (**Supp. Fig. 3B**) in shK8 cells, supporting BFA-related results, i.e., an implication of K8 in the ERAD pathway.

### 2. K8 is a critical factor for ERAD of μ_s_

To evaluate the potential role of K8 in ERAD, we took advantage of a cellular model system expressing the heavy chain μ_s_ subunit of secretory IgM. There, the lethal accumulation of μ_s_ in the ER due to the absence of the light chain to enable antibody reconstitution and secretion is counterbalanced by enhanced ERAD degradation. The implication of K8 in the degradation of μ_s_ was monitored via cell death upon K8-silencing (synthetic lethality assay, see Methods section for details).

K8 silencing triggered synthetic lethality of μ_s_-expressing cells (**Fig. 2A**; siK8+μ_s_). None of the other conditions (siRNA against K8 alone (siK8–μ_s_)) and μ_s_ expression alone (ctrl+μ_s_) led to significant cells death, asserting the features of *bona fide* synthetic lethality (**Fig. 2A**). These results showed that decreasing K8 expression compromised the ERAD pathway and led to enhanced apoptosis of μ_s_-accumulating cells.

**Figure 2.**
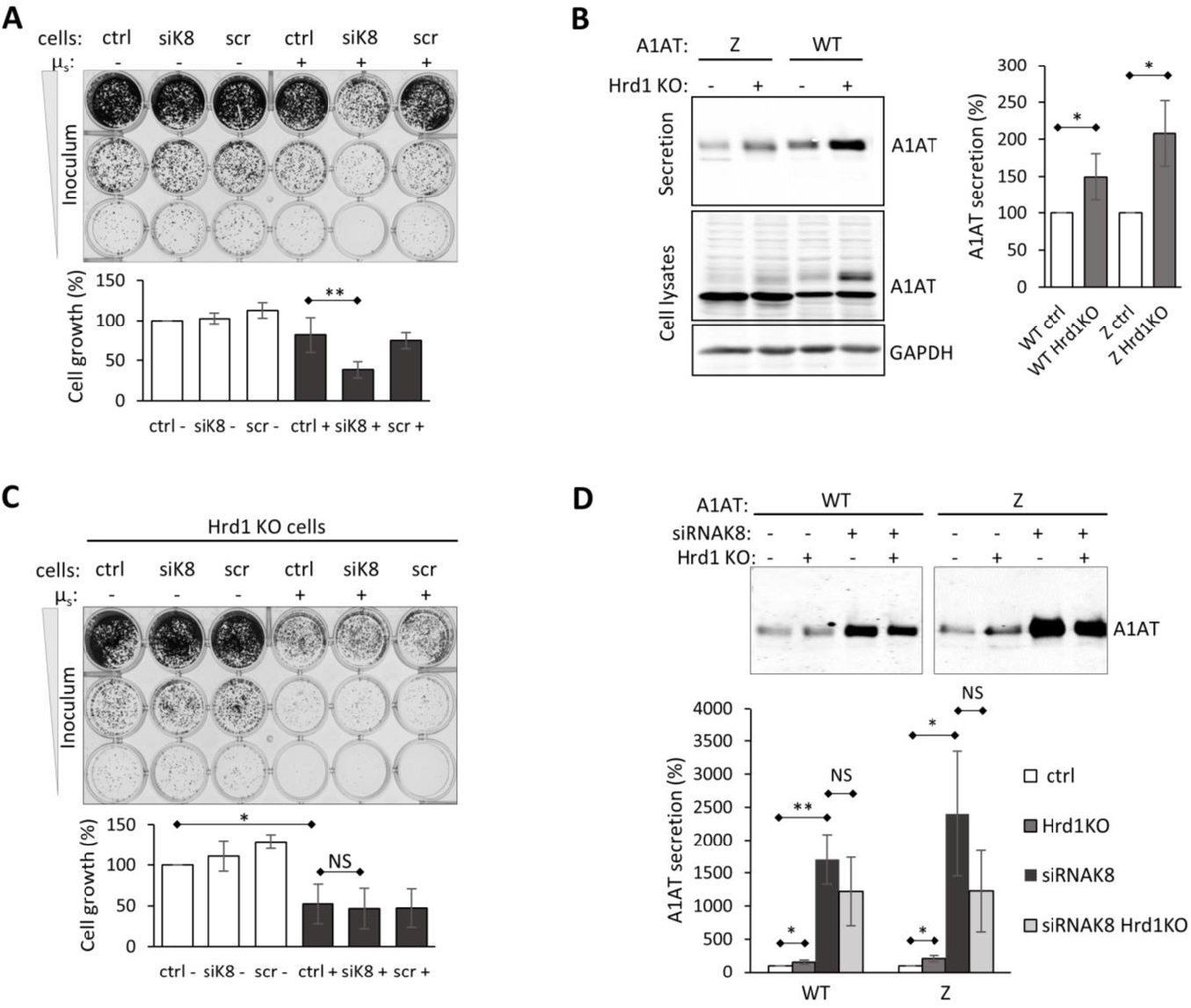
K8 regulates Hrd1-dependent processes **(A)** Synthetic lethality of HeLa cells conditionally expressing μ_s_. Cells transfected with siRNA against K8 (cells siK8), or scrambled siRNA (cells scr) were treated with 0.5 nM Mifepristone (Mif) (μ_s_+) or vehicle (μ_s_-). After transfection cells were seeded upon 1:5 serial dilution (inoculum 5,000, 1,000, 200 cells) into 24-well plates and grown for 7 days. Scrambled RNA had no effect on cell growth of non-expressing (scr-μ_s_) and μ_s_-expressing cells (scr+μ_s_). This is a representative image of a culture plate and quantification of cell growth (inoculum 1,000 cells, middle row) as mean % (±SD) of growth in non-transfected non-treated cells (ctrl-) of five independent experiments are shown. ** p< 0.005 (paired t test). **(B)** Effect of Hrd1 knock-out on WT-A1AT and Z-A1AT secretion from HeLa cells. WB images show WT-A1AT and Z-A1AT secretion levels (upper panel), A1AT expression (middle panel) and GAPDH as loading control (lower panel). Protein quantification is expressed as means ±SD from four independent experiments. * p < 0.05 (Mann-Whitney test). **(C)** Synthetic lethality of Hrd1KO HeLa cells. Cells were treated as in Fig. 2A. Representative image of culture plate and quantification of cell growth (inoculum 1,000 cells, middle row) as mean % (± SD) of growth in non-transfected non-treated cells (ctrl-) of five independent experiments are shown. * p < 0.05 (paired t test). **(D)** Effects of K8 silencing, Hrd1 knock-out and their combination on secretion of WT-A1AT and Z-A1AT. Upper panel: Normal (Hrd1KO-) and Hrd1KO (Hrd1KO+) HeLa cells were transiently transfected with WT-A1AT or Z-A1AT and with siRNA against K8 (siRNAK8+). Representative WB images demonstrating WT-A1AT and Z-A1AT secretion from normal, Hrd1KO, siRNAK8 and siRNAK8/Hrd1KO HeLa cells. Lower panel: Quantification of A1AT secretion as mean ± SD for three experiments. * p < 0.05, ** p < 0.005 (Mann-Whitney test).

### 3. K8 regulates Hrd1-dependent processes

We then searched for the ERAD process affected by K8 silencing. It has been previously shown that misfolded Z-A1AT secretion is not increased by inhibiting the proteasome (Teckman et al., 2001). We thus focused on ERAD steps governed by Hrd1 and p97/VCP (**Supp. Fig. 1**, Modules 2B and 3).

Knocking out Hrd1 in HeLa cells (stable Hrd1KO HeLa cells) (Vitale et al., 2019) increased Z-A1AT and WT-A1AT secretion (**Fig. 2B**), as already shown by Joly et al. (2017) for Z-A1AT. It was accompanied by increased levels of fully-glycosylated Z- and WT-A1AT in Hrd1KO cells (**Fig. 2B**). Therefore, disruption of the Hrd1 complex and consequently inhibition of substrate ubiquitination/retrotranslocation (**Supp. Fig. 1**, module 2A and 2B) appears to be an essential step favouring both WT-A1AT and Z-A1AT secretion.

To further investigate this question, we examined whether K8 is involved in the ERAD processes controlled by Hrd1. The effect of combining Hrd1KO and siRNAK8 was evaluated using synthetic lethality and A1AT secretion assays. Hrd1KO cells showed a significant synthetic lethality of cells upon μ_s_ induction (ctrl, siRNAK8, and scramble), underlining the importance of Hrd1 (**Fig. 2C**). When combining Hrd1KO and siRNAK8, no additive or synergistic effect was observed in synthetic lethality (**Fig. 2C**) or increased WT-/Z-A1AT secretion (**Fig. 2D**). Both growth and secretion assays showed that Hrd1 and K8 modulate the secretion of WT-/Z-A1AT *via* the same pathway.

Pharmacological inhibition of VCP/p97 protein with Eeyarestatin I or NMS-873 was performed to test for the role of substrate dislocation (**Supp. Fig. 1**, module 3) on WT-/Z-A1AT secretion (**Supp. Fig. 4**). Both inhibitors led to the intracellular accumulation of Z-A1AT and WT-A1AT, consistent with reduced degradation. Nonetheless, this was not associated with increased WT-/Z-A1AT secretion **(Supp. Fig 4** right top and bottom panels), indicating that the inhibition of substrate dislocation was insufficient to enhance secretion.

The observations above emphasised (i) the relationship between K8 and Hrd1 and (ii) the absence of increased secretion upon p97 inhibition, suggesting that K8 modulates early ERAD processes, i.e., Hrd1-dependent ubiquitination and initiation of retrotranslocation (**Supp. Fig. 1**).

### 4. K8 is recruited to ERAD complexes

For further insight into the involvement of K8 into ERAD processes, we determined if K8 is recruited to the ERAD complexes by performing sucrose gradient fractionations of ERAD complexes. The distribution of K8 and three important proteins of the Hrd1 complex (Hrd1, Sel1 and Derlin 2; **Supp. Fig. 1**, modules 2A and B) was monitored in mock-transfected cells and cells expressing WT-, Z-A1AT or μ_s_.

In mock-transfected cells, K8 was primarily associated with lower sedimentation fractions (3-6) together with Derlin2 and a smaller amount of Sel1 **(Fig. 3A-C**). This suggested that K8 forms complexes, at the least, with Derlin2 and Sel1 (**Fig. 7**).

**Figure 3.**
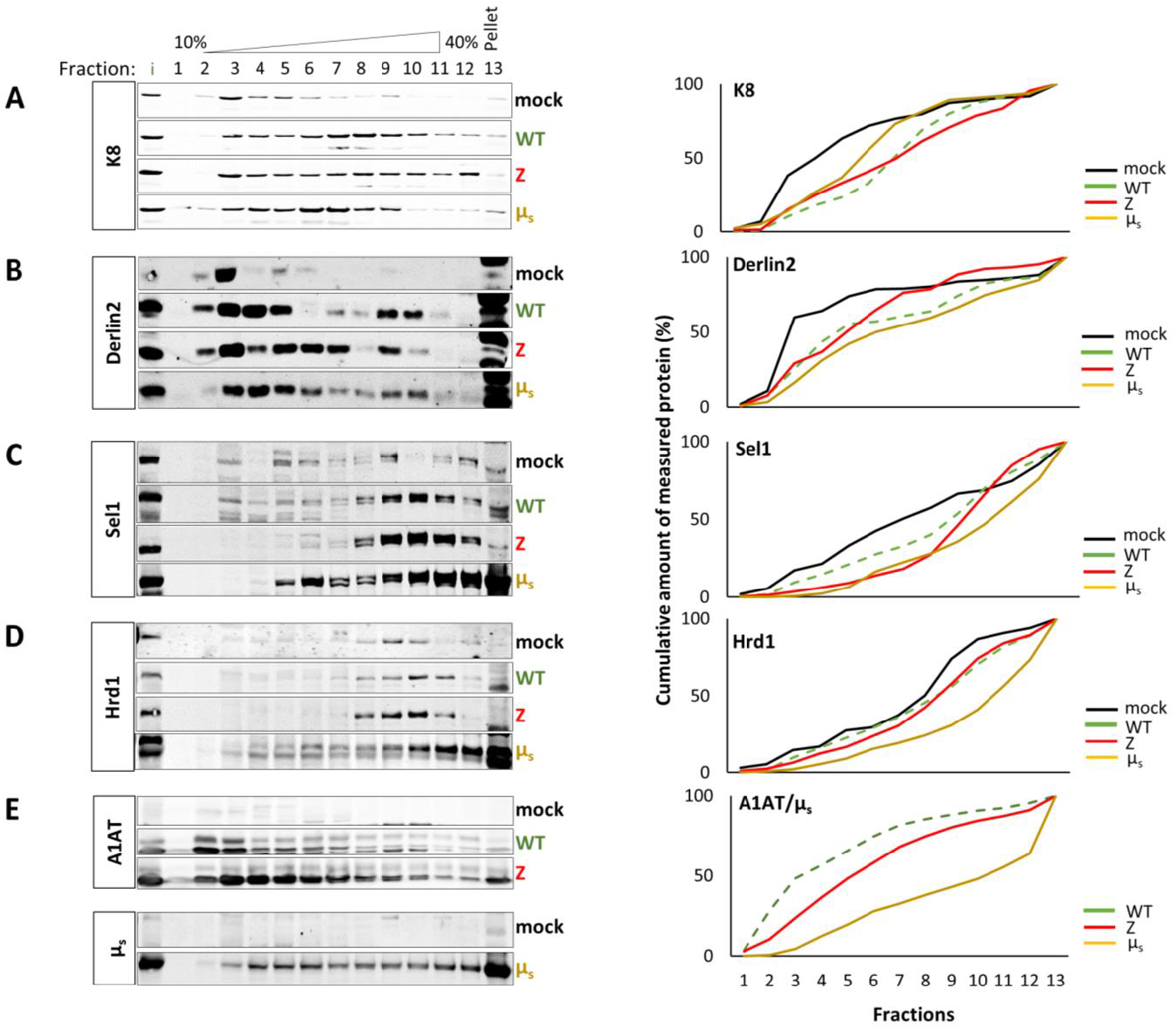
K8 is recruited to the ERAD complexes upon overexpression of misfolded Z-A1AT, WT-A1AT and μ_s_.Sucrose gradient fractionation of the ERAD complexes. HeLa cells expressing WT-A1AT (WT-A1AT), Z-A1AT (Z-A1AT), μ_s_ (Mif 0.5 nM for 24 h) (μs), or mock-transfected (mock). Protein samples from cells were sedimented over a 10–40% sucrose gradient. K8 (A), Sel1 (B), Hrd1 (C), Derlin-2 (D), and μ_s_/WT-A1AT/Z-A1AT (E) were detected by immunoblotting with respective antibodies and semi-quantified (right panels). Quantifications represent results shown in the corresponding images on the left and are expressed as cumulative amount of measured protein (%) to demonstrate the overall shift. In the left panel: black lines correspond to protein content in low density ERAD complexes under basal conditions, green, red, and yellow lines show ERAD complexes formed upon WT-A1AT, Z-A1AT and μ_s_ expression, respectively. The three tested substrates distributed differently in sucrose gradients: a large amount of μ_s_ which is a substrate efficiently processed by ERAD strongly distributed to heavy fractions, a relatively large amount of Z-A1AT distributed to heavy fractions, whereas smaller amounts of WT-A1AT were recruited to heavy fractions.

Expression of either Z-A1AT, μ_s_, or WT-A1AT, shifted K8 (**Fig. 3A**) together with Derlin2, Sel1 and Hrd1 (**Fig. 3B-D**) toward heavier fractions compared to mock-transfected cells. This is consistent with the dynamic recruitment of K8 to different higher-order ERAD complexes upon expression of an ERAD substrate (**Fig. 7**). The observed differences in the distribution of WT-A1AT, Z-A1AT and μ_s_ proteins in sucrose gradients may be due to the differences in the recruitment of these proteins to degradation process (**Fig. 3E**).

To test for the implication of K8 in the formation of ERAD complexes, we performed fractionation experiments on proteins obtained from shK8 mock- and shK8 Z-A1AT-transfected cells (**Fig. 4**). Under shK8 conditions, the total residual K8 co-sedimented with Derlin2 to one low-density fraction, unmasking a possible Derlin2-K8 complex (**Fig. 4A and B**, shK8 lines for K8 and Derlin2, black arrows). Profiles of Hrd1 and Sel1 fractionation were similar in mock-transfected and shK8 cells (**Fig. 4C and D**, compare lines shK8 and mock for Hrd1 and Sel1), suggesting that K8 does not influence the formation of complexes with Hrd1 and Sel1 under these conditions. The importance of K8 for the recruitment of Hrd1 and Sel1 to higher density complexes appeared on fractionation profiles of proteins derived from Z-A1AT expressing cells with normal and decreased levels of K8 (**Fig. 4C and D**).Noteworthy was that Hrd1 and Sel1 amounts were reduced in fraction 12 of the shK8 Z-A1AT sedimentation profile compared to cells expressing normal levels of K8 (**Fig. 4C and D**, red boxes). This shows that K8 favoured the recruitment of Hrd1 and Sel1 to high-order complexes under the expression of Z-A1AT.

**Figure 4.**
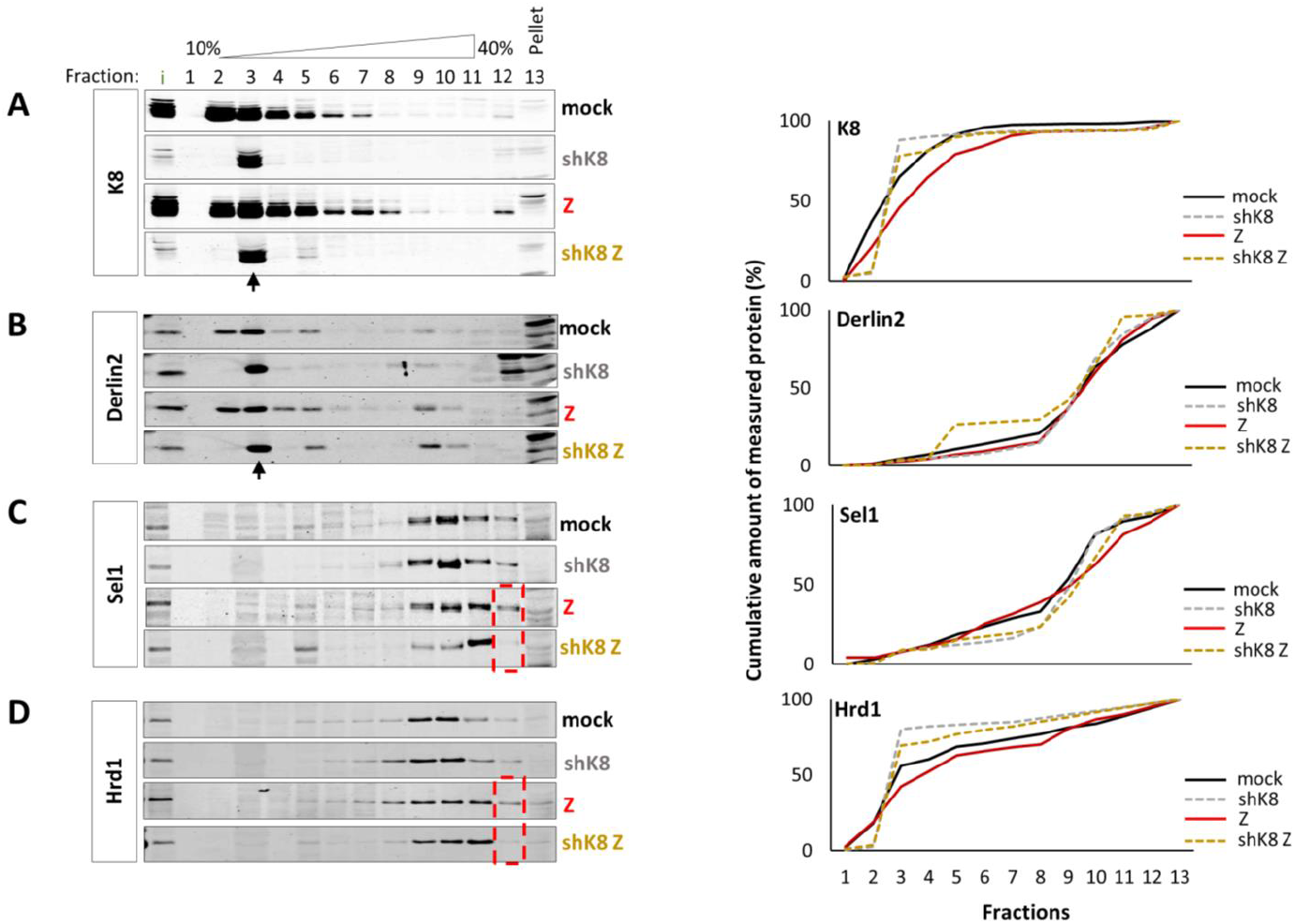
Silencing K8 affects ERAD complexes formation. Sucrose gradient fractionation of the ERAD complexes. HeLa cells expressing normal (mock and Z-A1AT) and decreased level of K8 (shK8 and shK8 Z-A1AT). Protein samples from cells were sedimented over a 10 - 40% sucrose gradient. K8 (A), Sel1 (B), Hrd1 (C),Derlin-2 (D) were detected by immunoblotting with respective antibodies. Black arrows correspond to ERAD complexes containing K8 and Derlin2 formed in shK8 cells, red dashed lined boxes correspond to high-density ERAD complexes formed in presence of K8 and Z-A1AT (Z-A1AT cells) but missing in shK8 Z-A1AT cells. Quantifications represent results shown in the corresponding images on the left and are expressed as cumulative amount of measured protein (%) to demonstrate the overall shift.

Co-staining of K8 and Calnexin (CNX) showed a redistribution of K8 to the vicinity of the ER in Z-A1AT-expressing cells *vs*. WT-A1AT and mock-transfected cells (**Supp. Fig. 5A**). K8 co-distributed with Z-A1AT in HeLa and primary HBE cells as shown by cytoimmunochemistry (**Supp. Fig. 5B and C**) and proximity ligation assay (**Supp. Fig. 5D and E**). Subcellular fractionation showed that K8 association with ER-containing microsomes (attested by the presence of Calreticulin, **Supp. Fig. 5G**) was enriched in Z-A1AT-expressing cells (**Supp. Fig. 5F**). Finally, Proteinase K assay experiments (Besingi and Clark, 2015) excluded intra-ER localisation of K8 (**Supp. Fig. 5G**). It was fully digested in both the presence and absence of detergent, as expected from the reported cytosolic localisation of K8 (according to Human Protein Atlas and (Coulombe and Wong, 2004)). Altogether, these observations are consistent with sucrose gradient fractionation results described above and support the implication of K8 in the degradation processes within ERAD.

To test whether other misfolded proteins recruited K8 to ERAD complexes, we conducted sucrose gradient fractionations of samples from WT-CFTR and F508del-CFTR-expressing 16HBE14o-cells. Analysis of protein complexes (**Fig. 5**) showed that Derlin2, Hrd1 and K8 shifted towards heavier fractions (**Fig. 5A – D**) upon F508del-CFTR expression compared to WT-CFTR, similar to that observed for Z-A1AT and μ_s_ (**Fig. 3A, B and D**). Noticeably, the Sel1 shift to heavier fractions was weaker on F508del-CFTR than Z-A1AT, WT-A1AT and μ_s_. The less pronounced shift of Sel1 towards high-density fractions upon F508del-CFTR expression could be explained because F508del-CFTR is expressed at an endogenous level compared to transiently transfected Z-A1AT.

**Figure 5.**
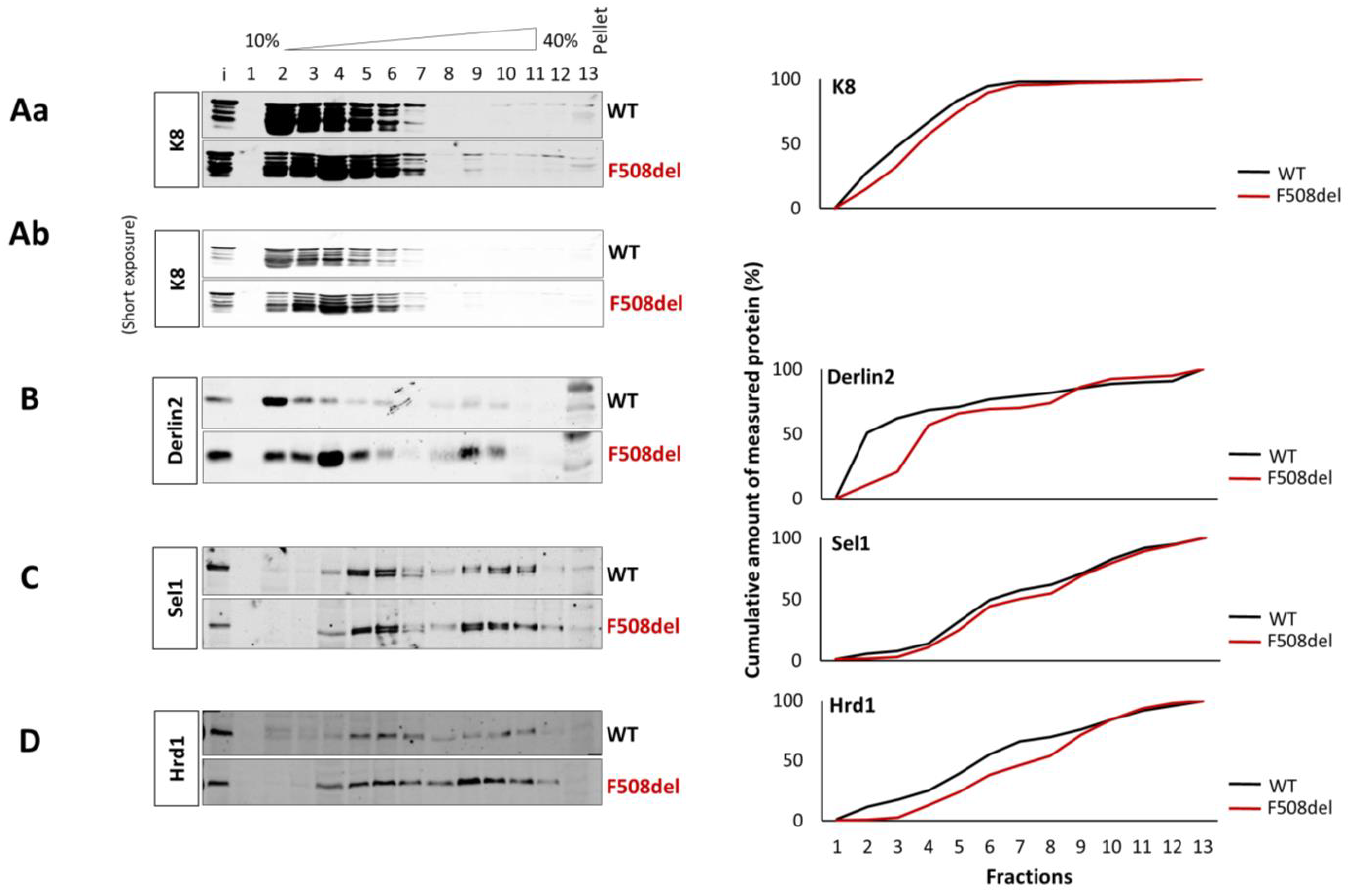
K8 is recruited to the ERAD complexes upon overexpression of misfolded F508del-CFTR. 16HBE cells expressing WT-CFTR and F508del-CFTR, as indicated, were lysed in 1% w/v LMNG and sedimented over a 10 - 40% w/v sucrose gradient. Levels of K8, Hrd1, Sel1, Derlin-2, were detected by immunoblotting. Representative images are shown. Quantifications represent results shown in the corresponding images on the left and are expressed as cumulative amount of measured protein (%) to demonstrate the overall shift.

Altogether, the results obtained for misfolded Z-A1AT and F508del-CFTR demonstrated that the delivery mechanism to degradation depends on K8 recruitment to the ERAD complexes (**Fig. 7**).

**5. A small molecule, c407, increases the secretion of Z-A1AT and is an ERAD modulator**

To test if K8-containing ERAD complexes were druggable, we measured the WT-A1AT/Z-A1AT secretion upon treating HeLa and primary human nasal epithelial (HNE) cells with the c407 compound. Treatment of both cell types resulted in an increase of Z-A1AT secretion in the concentration range of 10-35 μM (**Fig. 6A and B**), 20 μM c407 being the most efficient. In both primary HNE and HeLa cells, c407 did not affect WT-A1AT secretion.

**Figure 6.**
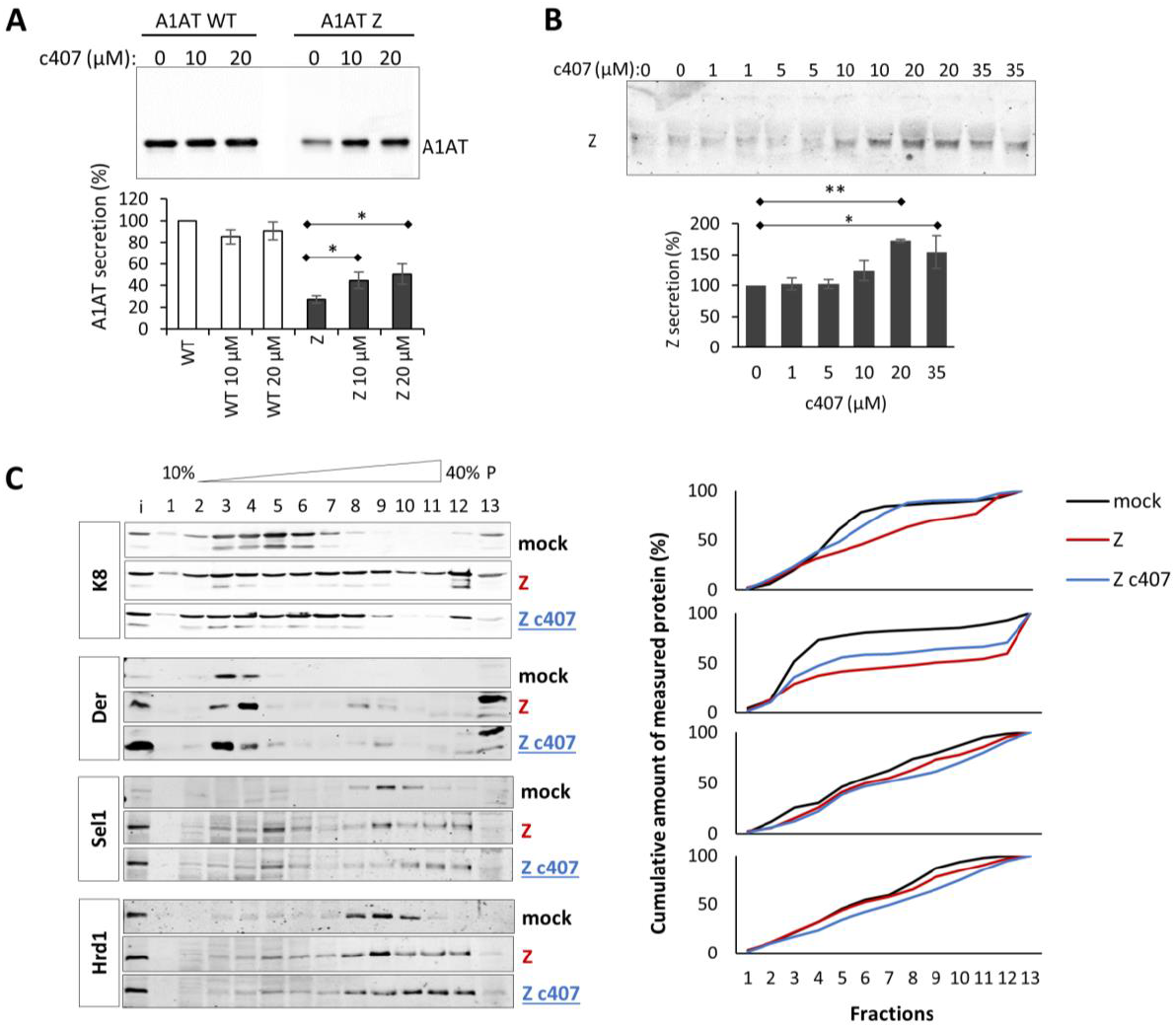
Compound c407 increases secretion of Z-A1AT from HeLa and primary HNE cells through modulating K8-contianing ERAD complexes. **(A)** Quantification of WT-A1AT and Z-A1AT secretion level from HeLa cells upon treatment with c407 (10 and 20) and vehicle (0). Cells were treated with c407 at concentrations of 10 and 20 μM for 24 h. The representative WB analysis image shows A1AT secreted into the medium. **(B)** Quantification of Z-A1AT secretion level from primary HNE cells upon treatment with c407 (1, 5, 10, 20 and 35) and vehicle (0). Cells were treated with c407 at concentrations of 1, 5, 10, 20 and 35 μM f or 14 days as pre-treatment and for 24 h for test of secretion. The representative WB analysis image shows A1AT secreted into the medium. **(C)** Sucrose gradient fractionation results upon c407 treatment. HeLa cells were transfected with Z-A1AT (Z) and treated with c407 at concentration of 20 μM or vehicle for 48 h, as indicated. Levels of K8, Hrd1, Sel1, and Derlin-2 were detected by immunoblotting in all fractions. Representative images are shown. Quantifications represent results shown in the corresponding images on the left and are expressed as cumulative amount of measured protein (%) to demonstrate the overall shift.

To test whether c407 targeted K8 directly, structure stabilisation experiments were performed using HDex-MS on purified K8. The addition of c407 (**Supp. Fig. 6A and B**) induced the stabilisation of multiple regions of K8, suggesting c407-K8 direct interactions. In solution, K8 was shown to exist as dimers in two distinct forms - a folded dimeric form stabilised by head-rod domain interactions and an unfolded dimeric form, as demonstrated by HDex-MS analysis (Premchandar et al., 2015). The robust allosteric stabilisation observed in the K8 head domain peptide 2-19 (**Supp. Fig 6A**, first panel) and the rod domain peptides (**Supp. Fig. 6A and B**) suggest a likely shift in the equilibrium to the folded form of the K8 dimer. Sucrose gradient fractionation of c407 treated HeLa cell lysates showed a less important shift of K8 and Derlin2 to heavy fractions compared to untreated cells (**Fig. 6C**), while Hrd1 and Sel1 were not affected. Altogether, these results suggest that targeting the K8 - Derlin 2 complex with c407 enhances Z-A1AT secretion.

## Discussion

In this study, we shed light on a new role of K8 as a scaffolding factor of ERAD protein complexes in epithelial cells (**Fig. 7**). We demonstrated that K8 negatively regulates secretory trafficking of misfolded Z-A1AT, as it was the case for F508del-CFTR (Colas et al., 2012a). Consequently, decreased K8 expression compromised ERAD leading to enhanced apoptosis of heavy chain μ_s_-expressing cells. Finally, we provided evidence of K8 implication in the ERAD processes by demonstrating K8 recruitment to complexes containing Derlin2, Hrd1 and Sel1 proteins and thus modulation of Hrd1-orchestrated processes (**Fig. 7**). Altogether, these data suggest that K8 participates in retrotranslocation initiation and ubiquitination of Z-A1AT, F508del-CFTR, μ_s_ and to some extent WT-A1AT and WT-CFTR. This unveils new structural and regulatory features of K8 within ERAD.

**Figure 7.**
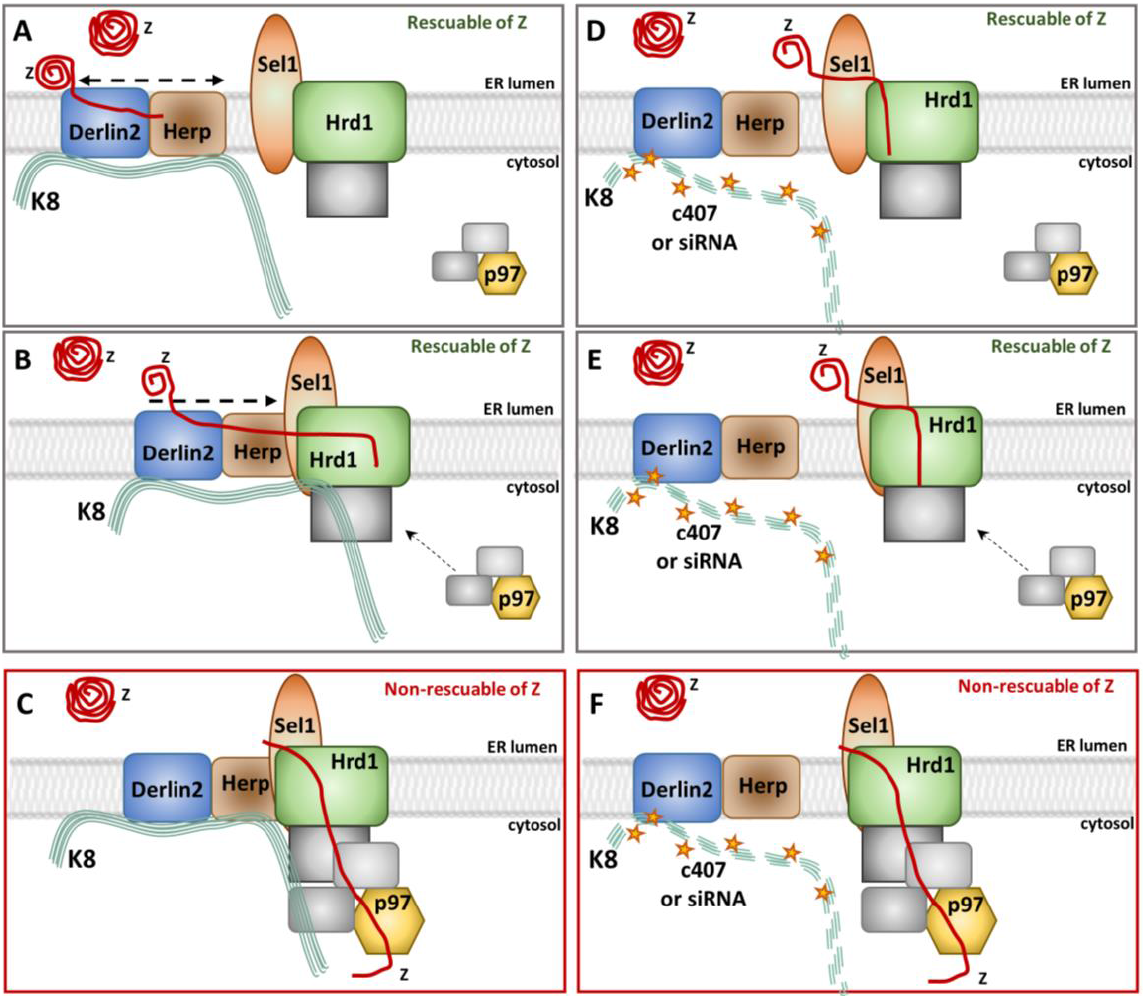
Model of K8 recruitment to the ERAD complexes at the ER membrane - hypothesis. (Adapted from Christianson and Ye). Normal K8 recruitment to the ERAD complexes: **(A)** K8 as a scaffolding platform in constant complex with Derlin2. Z-A1AT is recognised and recruited for retrotranslocation and ubiquitination. Stage rescuable for Z-A1AT secretion (upon Hrd1 or K8 down-regulation). Low density fractions complexes containing Derlin2 (module 2A in **Supp. Fig. 1**). **(B)** K8 facilitates complexing with Hrd1/Sel1. Retrotranslocation of ubiquitinated Z-A1AT is initiated. Stage rescuable for Z-A1AT secretion (upon Hrd1 or K8 down-regulation). Medium to high-density fractions complexes containing Derlin2, Sel1 and Hrd1 (modules 2A+2B in **Supp. Fig. 1**). **(C)** Z-A1AT is being dislocated from the ER lumen to the cytosol. Dislocation stage no rescuable for Z-A1AT secretion (upon p97 inhibition). Hypothetic higher-order complexes containing Derlin2, Sel1, Hrd1 and p97. Decreased or no recruitment of K8 to the ERAD complexes upon c407 molecule treatment or siRNA against K8: **(D)** K8-Derlin2 complex maintained (low-density fractions) even with residual K8. However, K8-dependent scaffolding of higher-order complexes is affected. Recruitment of Z-A1AT for retrotranslocation and ubiquitination is reduced. ER-luminal Z-A1AT is secreted. **(E)** K8-Derlin2 is not recruited to form complexes with Hrd1/Sel1. The shRNAK8 decreases Sel1 and Hrd1 recruitment to high-order complexes. By contrast, c407 treatment of cells do not change Hrd1 and Sel1 recruitment to higher-order complexes. Retrotranslocation initiation and ubiquitination of Z-A1AT is reduced. ER-luminal Z-A1AT is secreted. **(F)** Z-A1AT dislocation to the cytosol is reduced. ER-luminal Z-A1AT is secreted.

Misfolded Z-A1AT and F508del-CFTR are in soluble complexes with multiple ER chaperones (Schmidt and Perlmutter, 2005) (Kim and Skach, 2012). They are substrates for the ERAD pathway (Baldridge and Rapoport, 2016) (Vasic et al., 2020) (El Khouri et al., 2013) orchestrated by Hrd1-dependent ubiquitination at an early step (Wang et al., 2011) (Ramachandran et al., 2016). We showed agreement with previously published results (Joly et al., 2017) that Hrd1 knock-out led to the rescue of Z-A1AT secretion. K8 probably acts at this early step of ERAD, as it is recruited to the Hrd1-Derlin2 complexes. Furthermore, knockingout Hrd1, but not inhibiting p97, led to the rescue of the A1AT secretion. Altogether, these results suggested that reducing K8 expression diminishes ERAD efficacy, increases Z-A1AT maturation and, consequently, enhances Z-A1AT secretion. We postulate that K8 is a novel actor regulating ERAD. Furthermore, retrotranslocation is a crucial checkpoint defining rescuable and non-rescuable forms of misfolded proteins. This limit could be between Derlin2-Hrd1- and p97-governed steps of ERAD degradation when the retro-translocated substrate is undergoing dislocation (**Fig. 7 A and B *vs* C**).

We propose that K8 serves as a scaffolding platform on which other ERAD proteins build up functional complexes (**Fig. 7**). Different phases of retrotranslocation and ubiquitination are highly dynamic and create multiple stable and transient complexes upon stimuli (Eura et al., 2020). Therefore, it is not surprising that K8 can be recruited to many of them (**Fig. 7**) to facilitate transitions and remodelling of these complexes. The precise organisation of the retrotranslocating complexes is still not known. However, the involvement of the Hrd1-Hrd3 (Sel1) complex (Schoebel et al., 2017), Derlin family members (Ye et al., 2004) (Lilley and Ploegh, 2004), and the Hrd1-Der1 complex (Wu et al., 2020) has been shown. Importantly, our results suggest that (i) K8 – Derlin2 complexing is relatively permanent (detected even in mock cells and in cells with reduced K8 expression) and (ii) the recruitment of other ERAD proteins (Hrd1 and Sel1) occurs upon expression of misfolded Z-A1AT.

It could be argued that K8, an abundant protein in epithelial cells, binds non-specifically to intracellular protein complexes. However, as shown by immunocytochemistry, the intracellular distribution of K8 differed between Z-A1AT *vs* WT-A1AT-expressing HeLa and patient-derived HBE cells. It has also been previously observed for patients’ derived HBE cells expressing F508del-CFTR or WT-CFTR (Colas et al., 2012b). Moreover, non-specific K8 recruitment to ERAD complexes would be expressed as K8 sedimentation into all fractions in the shK8 cell and in mock-transfected cell lysates, which is not the case.

We postulate that keratins are essential regulators of processes occurring at the endoplasmic reticulum (ER) and Golgi. K18, which forms a heterodimer with K8, plays a role in F508delCFTR trafficking (Davezac et al., 2004) possibly by acting at the ER level (Toivola et al., 2010). Furthermore, K8 specifically binds Hsp70 involved in the ERAD pathway (Liao et al., 1995) (Salas et al., 2016) and K1, belonging to the same family of proteins as K8, forms complexes responsible for the retention of N-acetylglucosaminyltransferase within the Golgi structures (Petrosyan et al., 2015).

A small-molecule c407, which disrupts the interaction between F508del-CFTR and K8 (Odolczyk et al., 2013), increased Z-A1AT secretion in both primary differentiated HNE cells derived from patients and Z-A1AT-expressing HeLa cells. Because c407 modified K8 and Derlin2 recruitment to ERAD complexes, we propose that K8-Derlin2 complexes represent a potent target for pharmacotherapy. However, c407 treatment did not completely disrupt ERAD complexes because Sel1 and Hrd1 profiles did not change. Such an effect, if it occurred, would be toxic for cells and therefore inappropriate for therapy. C407 also failed to increase WT-A1AT secretion, contrary to reduced K8 expression, indicative of incompletely overlapping mechanism of actions which still need further studies to decipher. Using HDex-MS analysis, we showed that c407 stabilises multiple regions of K8 consistent with direct binding. HDex-MS experiments on Z-A1AT could not be performed due to its self-polymerisation.

Targeting the early steps of ERAD enables overcoming the difficulties of binding to highly mobile folding intermediates of Z-A1AT (Lomas et al., 2021). One of the early ERAD proteins, Hrd1, has been proposed as a direct therapeutic target for several diseases: rheumatoid arthritis (YAGISHITA et al., 2012) (Rahmati et al., 2018), type 2 diabetes (Wu et al., 2020), Alzheimer’s and Parkinson’s diseases (Nomura et al., 2016). Pharmacological or genetic inhibition of Hrd1 significantly reduced the severity of rheumatoid arthritis and glucose control in the diabetic model. By contrast, in Alzheimer’s and Parkinson’s diseases, Hrd1 overexpression leads to suppression of Pael-R-or Aβ1–40 and Aβ1–42-induced cell death (Nomura et al., 2016). Our therapeutical strategy, based on modulation of K8-regulated early ERAD complexes with chemical compounds opens new perspectives for A1ATD and other epithelial PMDs. Combining molecules targeting the misfolded protein directly (Lomas et al., 2021) with molecules affecting ERAD (e.g., c407) could lead to enhanced rescue.

In conclusion, we demonstrated that K8 is a scaffolding factor for early-stage ERAD complexes regulating the degradation of ERAD substrates and propose targeting ERAD complexes containing K8 as an attractive strategy for the pharmacotherapy of A1AD and possibly other epithelial PMDs.

## Materials and methods

### Cell culture

#### HeLa cells

HeLa cells transiently transfected with pcDNA3-WT-A1AT or Z-A1AT were cultured in Dulbecco’s modified eagle medium (DMEM) and were supplemented with 10% v/v fetal calf serum (FCS), 2 mM L-glutamine, 100 μg/ml streptomycin and 100 units/ml penicillin in a humidified incubator at 37°C 5% v/v CO_2_. WT-A1AT/Z-A1AT cDNA was kindly provided by Prof. Eric Chevet (University of Bordeaux, France). Western blot (WB) analysis of WT-A1AT and Z-A1AT extracted from transiently transfected HeLa cells revealed that Z-A1AT-expressing cells produced less fully-glycosylated A1AT than cells transfected with WT-A1AT (**Supp. Fig. 7A, upper left panel**) concomitant with reduced secretion, reaching only 25% of WT secretion level (**Supp. Fig. 7A, bottom left panel and quantification**).

We generated HeLa cells that express reduced amounts of K8 using the shRNA approach (shK8 HeLa cells). WB quantification showed that shK8 HeLa cells expressed ∼17% of the normal K8 level; transient transfection with either WT-A1AT or Z-A1AT did not affect this level (**Supp. Fig. 7B**).

HeLa cell lines stably expressing the IgM heavy chain subunit μ_s_ under induction with Mifepristone (Mif, 0.5 nM), WT variant and Hrd1KO, were a kind gift from Dr Eelco van Anken (IRCCS Ospedale, San Raffaele, Italy). These cells were cultured in DMEM (Dulbecco’s modified eagle medium) and were supplemented with 5% v/v FCS, 100 μg/ml streptomycin and 100 units/ml penicillin in a humidified incubator at 37°C and 5% v/v CO_2_.

#### Primary HBE/HNE cells

The establishment of primary bronchial epithelial cell cultures from PiZZ AATD patients (n = 3) and MM controls (n = 3) matched according to sex, GOLD stage (0 - III), and smoking status has been previously reported (van ’t Wout et al. 2014). Cell cultures were re-established from liquid nitrogen stored stocks. Primary nasal epithelial cell cultures from PiZZ AATD patients (n = 2) and MM control (n = 1) were established from fresh nasal brushing (Pranke et al., 2017). In primary human bronchial epithelial (HBE) and human nasal epithelial (HNE) cells derived from Z-homozygous (PiZZ) or WT-homozygous individuals (**Supp. Fig. 7C**), the fraction of secreted A1AT was lower (<50%) in PiZZ cells than in WT cells. In contrast, intracellular levels of Z-A1AT was increased compared to WT-A1AT (compare **Supp. Fig 7C upper and lower panels**).

### Silencing K8 expression

K8 expression was silenced in HeLa cells using a stable shRNA (Thermo Scientific) transduced by lentiviral vectors. HeLa cells containing the shRNA construct and parental HeLa cells were transfected with pcDNA3-A1AT WT or A1AT Z. For the growth or secretion assay in Hrd1KO cells, K8 expression was silenced with siRNA (Dharmacon siRNA Reagents) transiently transfected into HeLa cells with DharmaFect Reagent (Dharmacon).

### Protein sample preparation for immunoblotting

#### Cell lysis and cell extract preparation

Cells were first trypsinised and centrifuged at 300 g for 10 min at 4°C, washed on ice with phosphate-buffered saline (PBS) and recentrifuged at 300 g for 10 min at 4°C. The cell pellet was then resuspended in a lysis buffer composed of Tris-HCl pH 7.5, 20 mM, NaCl 50 mM, EDTA 1 mM, NP40 0.5% v/v and a protease inhibitor cocktail (cOmplete Tablets, Roche, 04 693 124 001) and incubated on ice for 30 min. Next, cell lysates were centrifuged at 1500 g for 15 min at 4°C. Supernatants were transferred to a new set of tubes, and the protein concentration was measured with the DC Protein Assay (BioRad, 500-0113, -0114, -0115) according to the manufacturer’s protocols.

#### Preparation of the secreted protein

Cells were grown to 80-90% confluence and then washed three times with PBS to remove serum. Cultures were then kept in the DMEM media containing only penicillin/streptomycin for 24 h or shorter. Media with secreted proteins were collected, centrifuged at 300 g, divided into aliquots and frozen at -80°C. Each sample used for the WB analysis was defrosted once.

### Western blot analysis

For protein detection by WB samples were mixed with a 5X Laemmli sample buffer and heated at 95°C for 5 min, resolved using a 10% w/v acrylamide SDS-PAGE gel and transferred to nitrocellulose membranes. The membranes were then incubated in a blocking buffer (3% w/v BSA (bovine serum albumin) and 3% w/v milk in PBS-0.1% v/v Tween-20) for 1 h. Proteins of interest were immuno-detected with appropriate antibodies (ab) diluted in a 3% w/v BSA blocking buffer for 1 h. K8 was immuno-detected with mouse monoclonal ab (61038, Progen) at a 1:500 dilution, A1AT was immuno-detected using the primary rabbit polyclonal anti-human A1AT ab (A0012, Dako, Agilent) at a 1:1,000 dilution, Derlin2 was detected with mouse monoclonal ab (sc-398573, Santa Cruz Biot Inc.). Sel1 was detected with mouse monoclonal ab (sc-377350, Santa Cruz Biot Inc.), and Hrd1 was detected with rabbit anti-Synoviolin ab (A302-946A—M, Bethyl). Other abs used were: rabbit polyclonal anti-GAPDH (sc-25778, Santa Cruz Biot Inc.) at a 1:500 dilution, mouse anti-Na^+^K^+^ATPase (ab7671, Abcam) at a 1:1,000 dilution, mouse monoclonal anti-Hsp70 (sc-24, Santa Cruz Biot Inc.) at a 1:1,000 dilution, rabbit anti-*Gaussia* luciferase (E8023, New England Biolabs) at a 1:2,000 dilution, rat monoclonal anti-Grp78 (sc-13539, Santa Cruz Biot Inc.) at a 1:500 dilution and rabbit polyclonal anti-Calreticulin (SPA-600, Stressgen) at a 1:500 dilution. Finally, nitrocellulose membranes were incubated with appropriate secondary antibodies coupled to fluorochromes, as recommended by the manufacturer (Li-Cor, Bad Homburg, Germany). The detection of the WB results was performed with the Odyssey scanner (Li-Cor, Germany). Quantification of the WB was performed using ImageJ software.

### Immunocytochemistry

HeLa and HBE cells were grown on microscopy slides up to low levels of confluence. The cells on the slides were washed twice with PBS and fixed with ice-cold acetone for 5 min. Next, the slides were washed twice with PBS, dried and stored at -20°C if necessary. The cells were then thawed, rehydrated with phosphate-buffered saline with 0.1% v/v Tween-20 (PBS-T) and incubated in blocking solution 3% w/v BSA in PBS-T for 1 h at room temperature (RT). Protein immunodetection was performed with primary abs diluted in blocking solution during overnight incubation at 4°C: rabbit polyclonal ab against alpha-1-antitrypsin (A0012, Dako, Agilent) at a 1:500 dilution, mouse monoclonal ab against K8 (61038, Progen) at a 1:200 dilution, and goat polyclonal ab against Calnexin (Santa Cruz Biot Inc.) at a 1:500 dilution. The primary antibodies against different proteins were incubated simultaneously. Subsequently, the cells were washed four times for 5 min each in PBS-Tween20 0.1% v/v, and nonspecific binding sites were blocked in 5% v/v goat serum (in PBS-Tween20) for 30 min. The cells were then incubated for 45 min at RT with goat secondary IgGs conjugated to Alexa 488 or 594 at a 1:1,000 dilution in 5% v/v goat serum in PBS-T. After five washes for 5 min each, the Vectashield mounting medium containing DAPI (Vector Laboratories, H-1200) was used to mount the cells on microscope slides. Immunocytochemistry on the air–liquid interface cultures of the primary HBE cells was performed, as described previously (Pranke et al. 2017).

### Confocal microscopy

Confocal microscopy was performed as described previously (Pranke et al., 2017). Briefly, a Leica TCS SP5 AOBS confocal microscope (Heidelberg, Germany) was used. Multiple optical z-stack images were captured over the cell culture. The images were analysed with ImageJ software (NIH, USA). The co-localisation level was assessed by the ImageJ plug-in JACoP and tools Image calculator and Measure stack. The PLA results were quantified with the Analyze Particles tool and expressed as an average number of fluorescent spots (nfs), minus the average number of negative controls.

### Growth assay

HeLa cells overexpressing IgM heavy chain subunits μ_s_ under Tet-inducible promoter (Bakunts et al. 2017) (Vitale et al. 2019) were kindly provided by Dr Eelco van Anken. In the absence of light chains, heavy chains cannot reconstitute IgM and cannot be secreted. The μ_s_ is retained in the ER, and its accumulation activates the unfolded protein response. Once the ERAD pathway is deactivated, heavy chains overload the ER leading to synthetic lethality through apoptosis, defining which factors are crucial to act in conjunction with Hrd1 and Sel1L in the disposal of μ_s_ (Vitale et al., 2019). The experiments were performed according to a previously published protocol (Bakunts et al., 2017). The HeLa cells overexpressing IgM heavy chain subunit μ_s_ upon induction with 0.5 nM Mifepristone (Mif) were first transfected with siRNA directed against K8 and scrambled sequence (Dharmacon siRNA Reagents). Non-transfected cells and cells 24 h after transfection were trypsinised, counted with a Malassez chamber, and seeded upon 1:5 serial dilutions (5,000, 1,000, and 200 cells per well) in 24-well plates. Mifepristone (0.5 nM) was added after cells were attached to the growth surface. To assess growth differences between tested conditions, cells were grown for 7 days. Culture media and pharmacological agents were refreshed every 2-3 days. Cells were then fixed with methanol-acetone (1:1) for 10 min, stained with 0.5% crystal violet in 20% methanol for 10 min, and washed with distilled water afterwards. Dried plates were imaged with the ChemiDoc TM XRS+ (BioRad). The intensity of crystal violet staining was quantified with ImageJ software (NIH). An average intensity of empty wells on the same plate served for background subtraction. Quantification of growth was presented for wells with a seeding of 1,000 cells.

### ERAD complexes fractionation

The experiments were performed as described in (Vitale et al. 2019). HeLa cells were grown on T75 flasks up to 75% of confluence and transfected with plasmids coding for WT-A1AT or Z-A1AT using the Lipofectamine 3000 protocol (Thermo Fisher Scientific). Additionally, heavy chains μ_s_ were induced with Mifepristone 0.5 nM 24h before harvesting the cells. Cell lysis was performed 48h post-transfection with a lysis buffer (50 mM Tris-HCl pH 7.4, 150 mM NaCl, 5 mM EDTA, 1% lauryl maltose neopentyl glycol (LMNG)) containing a mix of proteases inhibitors (cOmplete Tablets, Roche, 04 693 124 001). After centrifugation at 5,000 g for 10 min, cell lysates were loaded on the top of 10 % - 40 % linear sucrose gradients and centrifuged at 39,000 rpm / 17 h / 4°C in a SW41Ti rotor. Sucrose gradients were prepared with a Hoefer gradient maker. The 13 fractions were collected from low density at the top of the gradient to high density at the bottom. Proteins were precipitated with trichloroacetic acid and washed with ice-cold acetone. Protein pellets were resuspended in a Laemmli buffer containing dithiotreitol (DTT, 10 mM), heated at 50°C before separation by SDS-PAGE. After protein transfer to nitrocellulose membrane, proteins were detected with appropriate antibodies: K8 – mouse monoclonal ab (61038, Progen), Sel1 (sc-377350, Santa Cruz Biot Inc.), Hrd1/Synoviolin (A302-946A-M, Bethyl), and Derlin2 (sc-398573, Santa Cruz Biot Inc.). The intensity of each protein band was measured using ImageJ (NIH Bethesda) and expressed as % of a sum of all fractions.

### Preparation of cell microsomes

The cells grown in flasks were washed with PBS, trypsinised and centrifuged at 300 g for 5 min at 4°C. The cell pellet was washed twice in PBS, recentrifuged at 300 g for 5 min at 4°C and resuspended in 800 μl of the isotonic extraction buffer (10 mM HEPES, pH 7.8, 250 mM sucrose, 25 mM potassium chloride and 1 mM EDTA) in the presence of a protease inhibitor cocktail (Roche). Cell lysis was performed using mechanical homogenisation with constant incubation of the samples on ice. Up to 70% of the sample showed disrupted cells when observed under microscope. The homogenised cells were then transferred to prechilled 1.5 ml tubes and centrifuged at 6000 g for 10 min at 4°C; the collected supernatant was recentrifuged in a fresh tube at 9000 g for 15 min at 4°C. A new supernatant was collected in a prechilled 1.5 ml ultracentrifuge tube and centrifuged at 100000 g for 1 h at 4°C in a TLA-55 rotor (Beckman Coulter). The resulting pellet was resuspended in the same isotonic buffer containing antiproteases and was frozen at -80°C together with the 100000 g supernatant, which represented the cytosolic fraction.

### Proteinase K assay on the microsomal fraction

The Proteinase K assay, as adapted from Besingi and Clark (2015), was designed to detect the presence or absence of a K8 protein in the interior of the ER/microsomal compartment or at the cytosolic periphery of the ER membrane. HeLa cells transfected with cDNA A1AT were grown in a T150 culture flask, trypsinised, washed with PBS and pelleted at 300 g for 5 min at 4°C. The microsomal fraction was purified, as described in the section ‘Cell microsomes preparation’. The microsomes were resuspended in the isotonic buffer. Each of the samples containing the microsomes was divided into two volumes. The first half of the microsomes was diluted in the same isotonic buffer, and the second was diluted in the isotonic buffer containing 1% v/v Triton X-100 (final Triton X-100 concentration 0.5% v/v) and incubated for 15 min to solubilise membranes and release intramicrosomal proteins. Next, the two samples (with and without Triton X-100) were divided to retain a nondigested aliquot; Proteinase K was added to the second aliquot to a working concentration of 125 μg/ml. It is assumed that Proteinase K digests all proteins in Triton X-100-treated samples and only externally/transmembrane-located proteins in nontreated samples. All samples – (i) microsomes in isotonic buffer; (ii) microsomes in isotonic buff. + Proteinase K; (iii) microsomes + Triton X-100; (iv) microsomes + Triton X-100 + Proteinase K – were incubated at 37°C for 20 min. All samples were resolved on 10% SDS-PAGE gels and immunoblotted to detect A1AT (Dako, Agilent), K8 (Progen, Germany), Calreticulin (Stressgen) and Na^+^K^+^ATPase (Abcam).

### Cell culture treatments

HeLa cells treatments with the VCP/p97 inhibitor Eeyarestatine I (10 μM, Sigma Aldrich) in DMSO were done for 8 h, and NMS-873 (1μM, Selleckchem) in DMSO for 20 h, at 37°C in the CO_2_ incubator. HeLa cells were treated with 5μg/ml Brefeldin A for 7 h at 37°C in the CO_2_ incubator. Mifepristone was used at concentration of 0.5 nM for 6 days in the growth assay or for 48 h for secretion assay. Corrector molecule c407 was tested at final concentrations of 1, 5, 10, 20, and 35 μM.

### Statistics

The experiments were repeated at least three times. The results are expressed as mean ± SD and analysed using Mann-Whitney test or as described for specific experiments.

## Acknowledgements

The authors thank Dr. Eric Chevet for A1AT/Z-A1AT plasmids, Annemarie van Schadewijk for help in culturing the HBE cells, and Anush Bakhunts for μ_s_ expressing HeLa cells. We thank Dr Grazyna Faure, Dr. Stefano Fumagali and Dr Olivier Namy for helpful discussions and advices. We thank Muriel Girard and Dominique Debray for providing the A1AT/Z-A1AT primary cells. We are very grateful to Isabelle Hatin for technical assistance, and the cell imaging platform for assistance with microscopy experiments.

## Authors’ contributions

Iwona Maria Pranke participated in the experimental strategy preparation, conceived the protocols, performed experiments, analysed and interpreted results and wrote the manuscript; Benoit Chevalier performed biochemistry experiments; Aiswarya Premchandar performed mass spectrometry experiments and analysis; Nesrine Baatallah and Kamil F. Tomaszewski performed some biochemistry experiments; Sara Bitam established the shRNAK8 cell line; Danielle Tondelier performed cell biology experiments; Anita Golec participated in primary cell culture and biochemistry experiments; Jan Stolk provided the A1AT/Z-A1AT primary cells; Gergely L. Lukacs participated in the experimental strategy preparation; Pieter S Hiemstra, Michal Dadlez and David A Lomas participated in the writing of the manuscript; James A Irving participated in preparing the experimental strategy and edited the manuscript; Agnes Delaunay-Moisan participated in designing the experimental strategy and editing of the manuscript; Eelco van Anken participated in preparing the experimental strategy; Alexandre Hinzpeter performed cell biology experiments and wrote the manuscript; Isabelle Sermet-Gaudelus participated in preparing the experimental strategy and writing of the manuscript; Aleksander Edelman conceived of and coordinated the project and wrote the manuscript. All authors reviewed the manuscript and approved its submission.

