## Supplemental Figures for "Keratin 8 is a scaffolding and regulatory protein of ERAD complexes"

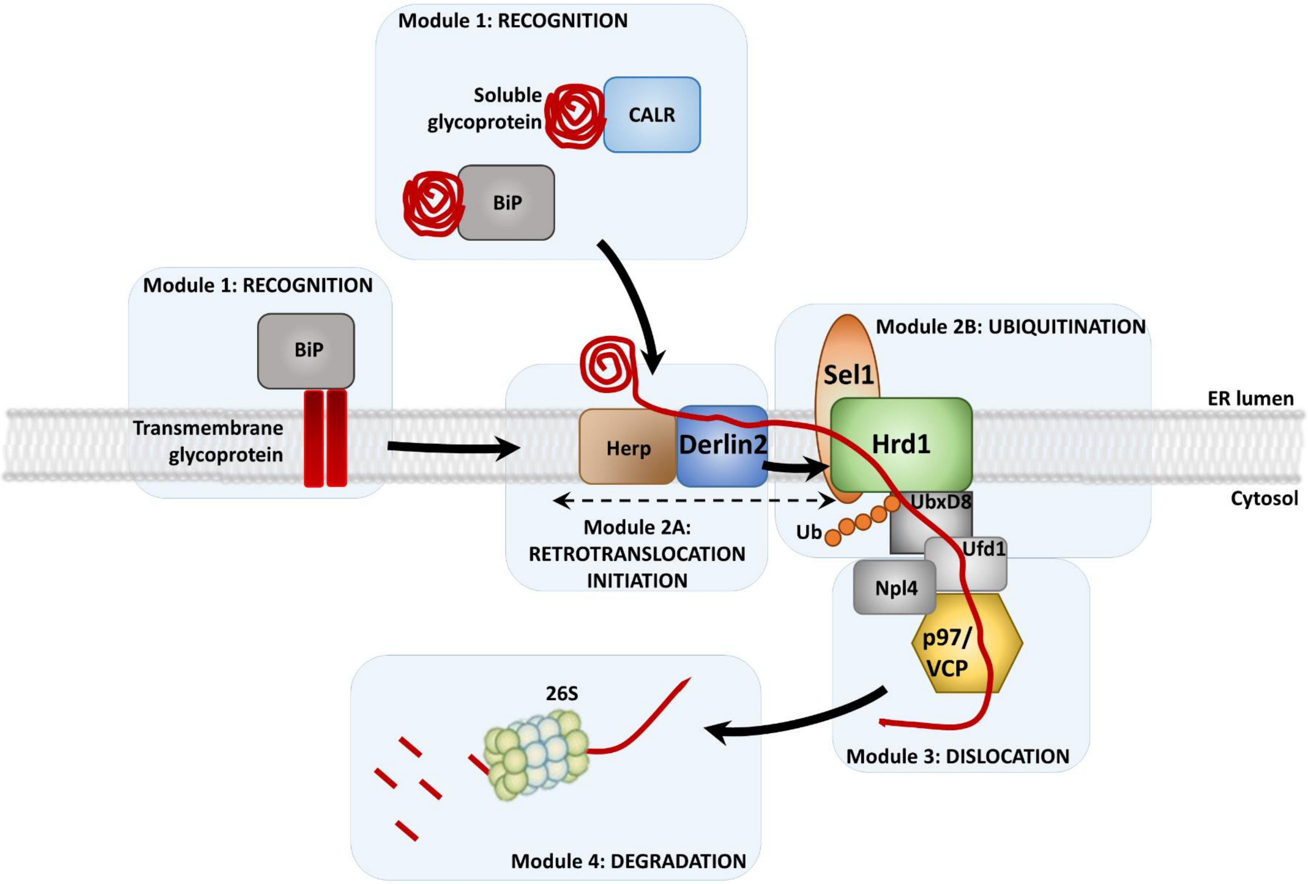


**Supplementary Figure 1.** Schematic representation of the mammalian ERAD pathway. **(Adapted from Christianson and Ye)**

Multiple ERAD proteins are organised into several modules. In general, proteins in the same module tend to form stable interactions, whereas proteins belonging to different modules dynamically bind with each other. Module 1 – Recognition: degradation substrates are bound to various chaperone proteins and glycosylation/deglycosylation factors, such as BiP and Calreticulin. Substrate recognition mechanisms vary depending on the type of substrate whether soluble or transmembrane and folded or misfolded. Module 2A – Retrotranslocation initiation: with substrate unfolding assured mainly by multifunctional Derlin proteins. Derlins participate in the transmembrane pore formation together with other proteins, i.e. Hrd1 E3 ubiquitin ligases. Module 2B – Ubiquitination: ubiquitin conjugation is mediated by E3 ubiquitin ligases including Hrd1 coupled to substrate recognition protein Sel1, and RNF185. Module 3 – Dislocation: extraction of the polypeptide from the ER membrane *via* the p97/VCP ATPase complex providing a pulling force. Dislocated polypeptides are targeted to the 26S proteasome for degradation (Module 4 – Degradation). The arrows indicate the flow of the substrates. The dashed line-arrow indicates the recruitment or dissociation of Derlin2 from complexes of module 2B.

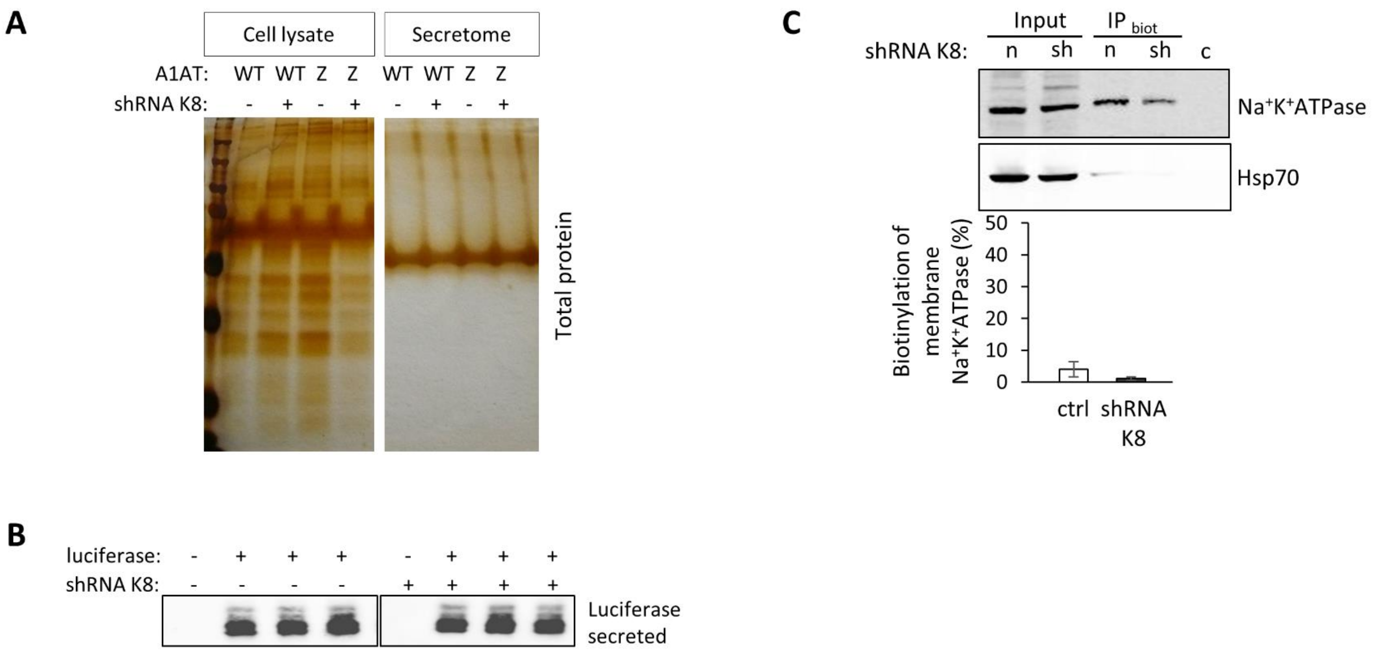


**Supplementary Figure 2.** K8 silencing does not influence global secretory trafficking.

**(A)** Silver-stained gel represents the cell lysates total protein content and the secretome protein content for shK8 and control HeLa cells.
**(B)** Transfected *Gaussia* luciferase secretion level from shK8 and control HeLa cells analysed using WB.
**(C)** Surface biotynilation of $Na^{+}K^{+}$ATPase in shK8 and control HeLa cells.

**A**

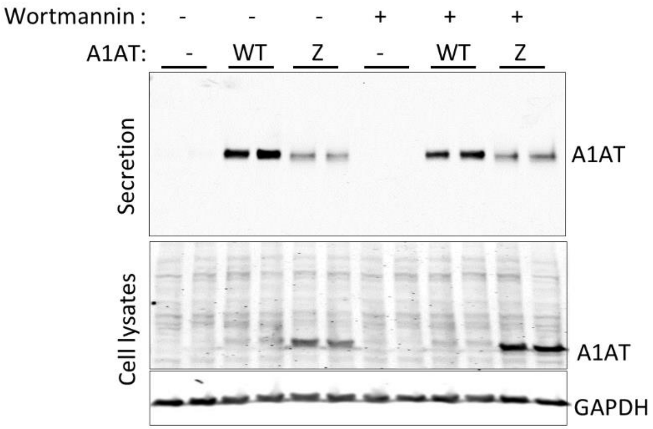


**B**

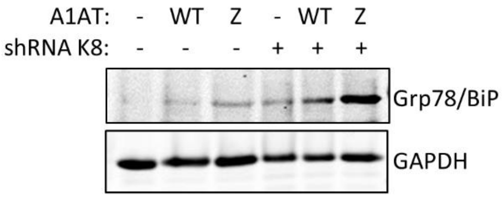


**Supplementary Figure 3.** K8 silencing increase ER stress in HeLa cells expressing Z-A1AT. Autophagy inhibition does not give the rescue of secretion.

**(A)** Immuno-analysis of secreted and intracellular WT- /Z-A1AT after HeLa cells treatment with Wortmannin (1 µM) for 24 h.
**(B)** Immuno-analysis of Grp78/BiP expression level in shK8 and control HeLa cells.

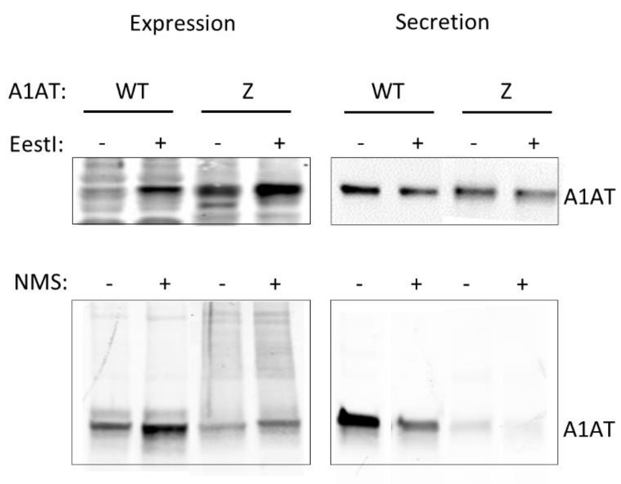


**Supplementary Figure 4. (Related to Figure 2)** Effect of p97 inhibition with EestI and NMS-873 inhibitors on A1AT secretion and accumulation in HeLa cells.

Representative WB immunodetection of WT- /Z-A1AT expression and secretion from HeLa cells in DMSO control conditions and upon treatment with EestI (10 µM) for 7 hours (top panels) and NMS-873 (1µM) for 20 hours (bottom panels).

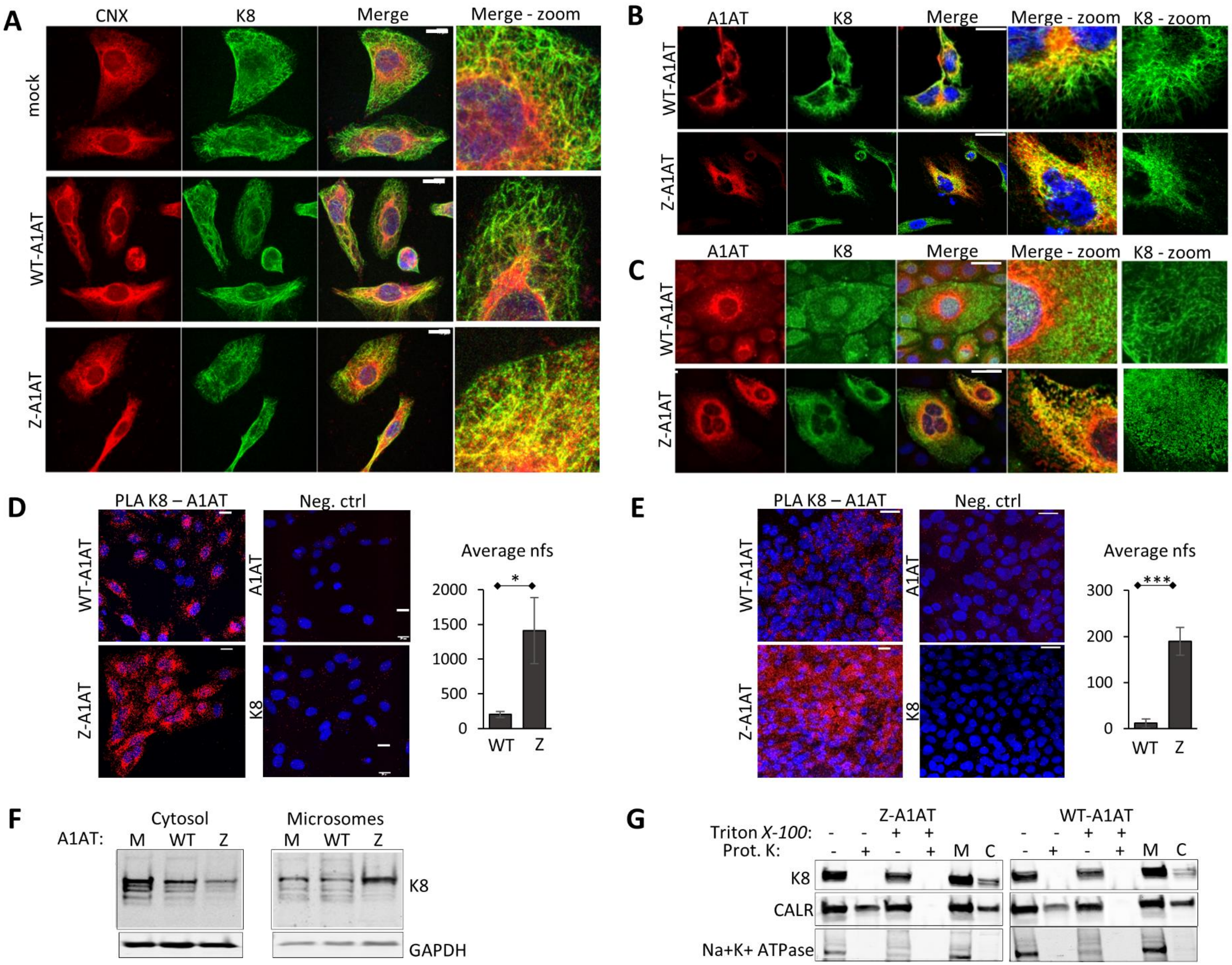
A
CNX
K8
Merge
Merge - zoom
mock
WT-A1AT
Z-A1AT
B
A1AT
K8
Merge
Merge - zoom
K8 - zoom
WT-A1AT
Z-A1AT
C
A1AT
K8
Merge
Merge - zoom
K8 - zoom
WT-A1AT
Z-A1AT
D
PLA K8 – A1AT
Neg. ctrl
WT-A1AT
Z-A1AT
A1AT
K8
Average nfs
*
2000
1500
1000
500
0
WT
Z
E
PLA K8 – A1AT
Neg. ctrl
WT-A1AT
Z-A1AT
A1AT
K8
Average nfs
***
300
200
100
0
WT
Z
F
A1AT:
Cytosol
M WT Z
Microsomes
M WT Z
K8
GAPDH
G
Z-A1AT
WT-A1AT
Triton X-100: - - + + - - + +
Prot. K: - + - + M C - + - + M C
K8
CALR
Na+K+ ATPase

**Supplementary Figure 5.** K8 redistributes to the ER upon expression of Z-A1AT in HeLa and primary HBE cells.

**(A)** Immunocytochemistry of A1AT and Calnexin (CNX), the ER marker, in mock-, WT-A1AT- or Z-A1AT-transfected HeLa cells. Representative images are shown. Scale bar = 20 µM. Right panel: Quantification of % of A1AT staining surface colocalising with K8 staining in HeLa cells (three independent experiments).

**(B)** Immunocytochemistry of A1AT and K8 in HeLa cells transfected with WT-A1AT or Z-A1AT. Representative images are shown. Scale bar = 20 µM.

**(C)** Immunocytochemistry of A1AT and K8 in the primary HBE cells from the WT-homozygous donor and Z-homozygous patient, grown on microscopy slides. Representative images are provided. Scale bar = 20 µM.

**(D)** Proximity ligation assay for K8 and A1AT in HeLa cells. Quantification represents an average number of fluorescent spots (nfs) / cell.

**(E)** Proximity ligation assay for K8 and A1AT in primary HBE cells. Quantification represents an average number of fluorescent spots (nfs) / cell.

**(F)** Immunodetection of K8 and GAPDH in cytosolic and microsomal fractions of mock- (M), WT-A1AT- (WT), and Z-A1AT-transfected (Z) HeLa cells. Representative images of WB detection.

**(G)** HeLa cells derived microsomes were treated with Proteinase K (125 µg/ml) alone and Triton X-100 (0.5%) followed by Proteinase K. Microsomes were prepared using homogenisation of HeLa cells in an isotonic buffer and sequential centrifugations. Half of the volume was pre-treated with Triton X-100 (0.5%) and both types of samples were incubated with Proteinase K (125 µg/ml) at 37°C. The levels of degradation of K8, ER Calreticulin and transmembrane $Na^+K^+$ATPase were evaluated using WB and compared to untreated samples with Proteinase K. Samples M ctrl – microsomes (pellet of 100,000 g; C ctrl – cytosolic fraction (supernatant 100,000 g). Representative results are shown for cells expressing WT-A1AT or Z-A1AT.

### Supplementary Figure 6 A

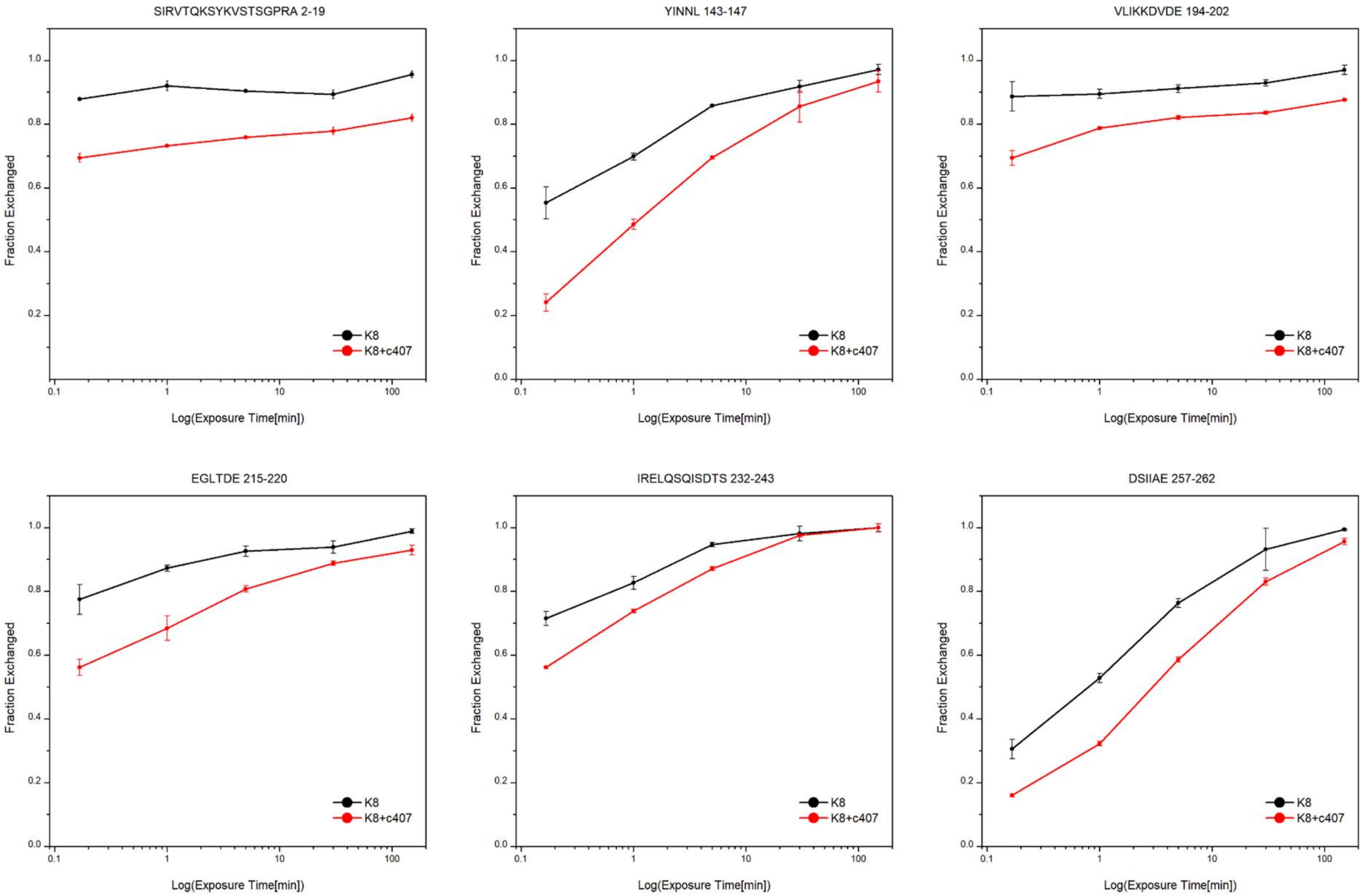

#### Supplementary Figure 6 B

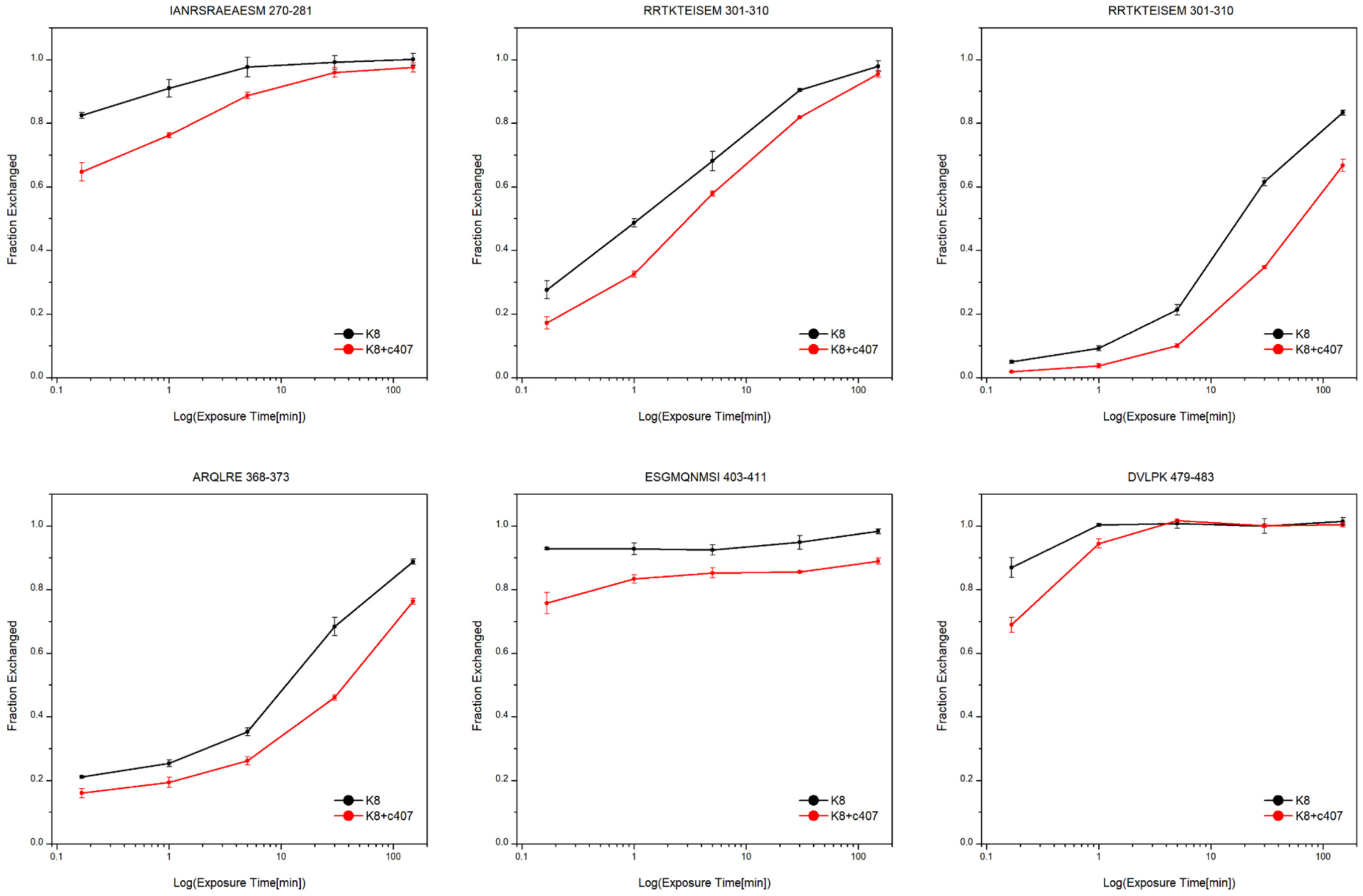

**Supplementary Figure 6 A and B.** Kinetic plots of all the identified peptides from K8, depicting the fraction exchanged as a function of time.

The unbound, apo state is shown in black, while the bound, K8-c407 complex is shown in red. Error bars represent the standard deviation of three independent experiments.

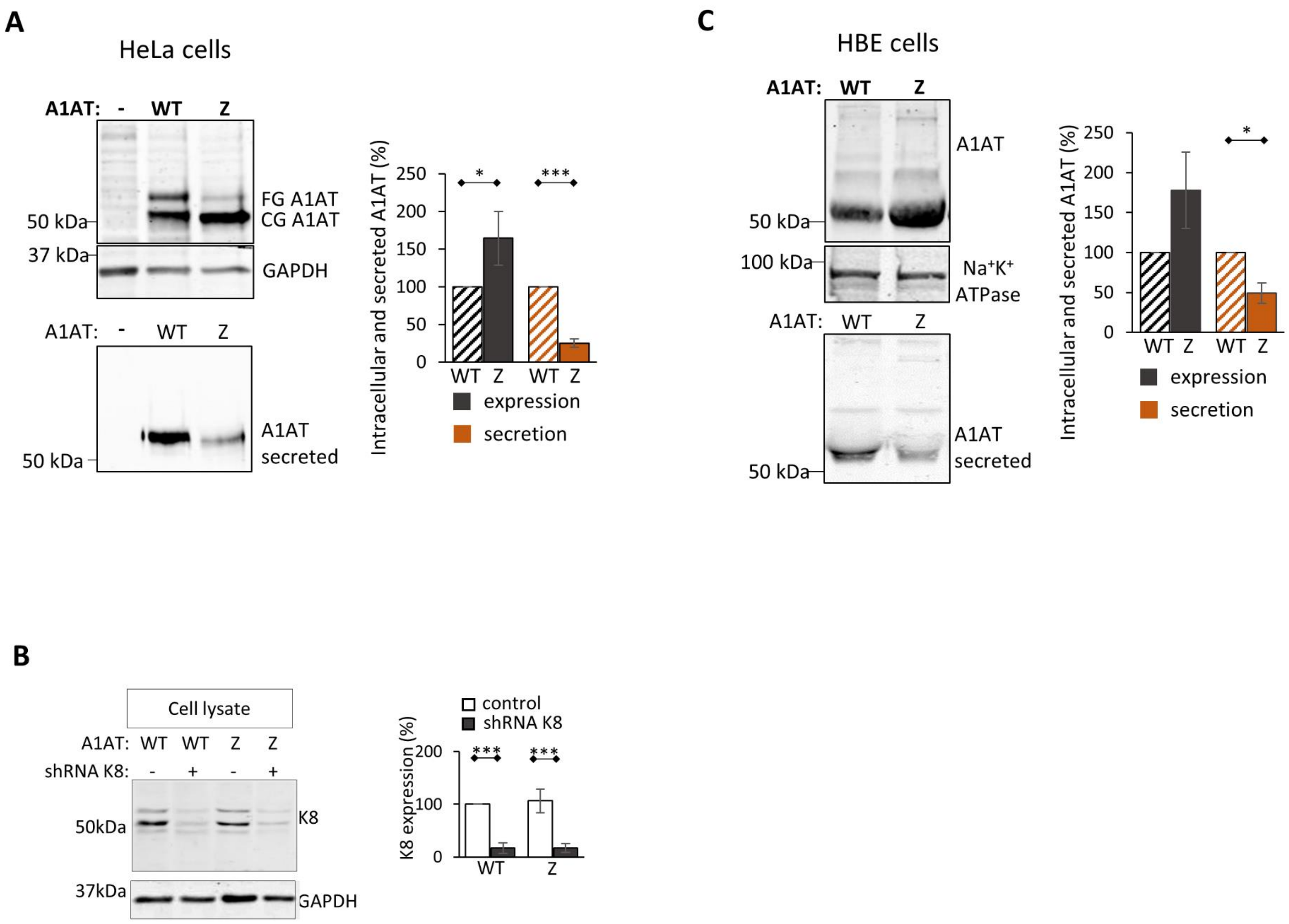


**Supplementary Figure 7. (Related to Material and methods)** Expression of WT-A1AT and Z-A1AT in HeLa and primary HBE cells – characterization of cells

**(A)** Expression and secretion of WT-A1AT and Z-A1AT in HeLa cell cultures after transfection with pcDNA3 plasmids coding for WT-A1AT or Z-A1AT. Top membranes - cell extracts probed for A1AT expression and GAPDH as a loading control. Bottom membrane – culture media without FCS after 24 h incubation with transfected HeLa cells were probed for the presence of A1AT. FG – fully-glycosylated, CG – core-glycosylated. Representative images are included. Quantification of nine independent experiments (* $p \leq 0.05$, *** $p \leq 0.0001$).

**(B)** Expression and secretion levels of WT-A1AT and Z-A1AT in primary HBE cell cultures isolated from two healthy controls (wt/wt A1AT) and two homozygous Z-A1AT individuals. Top membranes - cell extracts probed for expression of A1AT and $Na^{+}K^{+}$ATPase as the loading control. Bottom membrane - culture media without FCS after 24 h incubation with HBE cells were analysed for the presence of A1AT. Representative images are demonstrated. Quantification of four experiments for each subject (* $p \leq 0.05$).

(C) Western blot detection and quantification for intracellular K8 and GAPDH in cells with normal (shRNAK8-) and decreased (shRNAK8+) levels of K8 expression.
